# Mitochondrial dysfunction Impairs the Nuclear Pore Complex in Parkinson’s Disease Pathogenesis

**DOI:** 10.1101/2024.11.05.622143

**Authors:** Zainab Riaz, Yuan-Teng Chang, Gary Zenitsky, Huajun Jin, Vellareddy Anantharam, Arthi Kanthasamy, Anumantha Kanthasamy

**Affiliations:** Isakson Center for Neurological Disease Research, The University of Georgia, Athens, GA 30602; Department of Biology, The University of Georgia, Athens, GA, 30602; Department of Biochemistry and Molecular Biology, The University of Georgia, Athens, GA 30602; Department of Physiology and Pharmacology, The University of Georgia, Athens, GA 30602

## Abstract

The mislocalization of transcription factors and irregularities in the nuclear envelope of dopaminergic neurons affected by Parkinson’s disease (PD) implicates the nuclear pore in disease pathogenesis. While mitochondrial dysfunction is an integral component of PD pathophysiology, the involvement of channel-forming nucleoporins (Nups) in mitochondrial dysfunction-related neurodegeneration has not been investigated. Here we have identified pathological changes in the levels and distribution of a set of Nups, which are key structural and functional components of the nuclear pore complex, in dopaminergic neuronal models of PD. We observed that mitochondrial dysfunction reduces the expression of these Nups and disrupts the localization of Ran GTPase in both in vitro and in vivo dopaminergic neuron models. Furthermore, the nuclear pore central channel component, Nup62, mislocalizes and accumulates in the cytoplasm of these neurons under mitochondrial stress conditions. Mitochondrial stress also interferes with classical nuclear export of proteins in dopaminergic neural cells. Notably, we observed Nup pathology and Ran gradient loss in nigral dopaminergic neurons of PD patient brains, which highlights the clinical relevance of nuclear pore dysfunction as a disease mechanism. These findings provide direct evidence of the critical role of Nup-related abnormalities in mitochondrial dysfunction-induced degeneration of dopaminergic neurons, ultimately connecting nuclear pore complex and nucleocytoplasmic transport dysregulation to cell death in the development of PD.

**Graphical Abstract:** 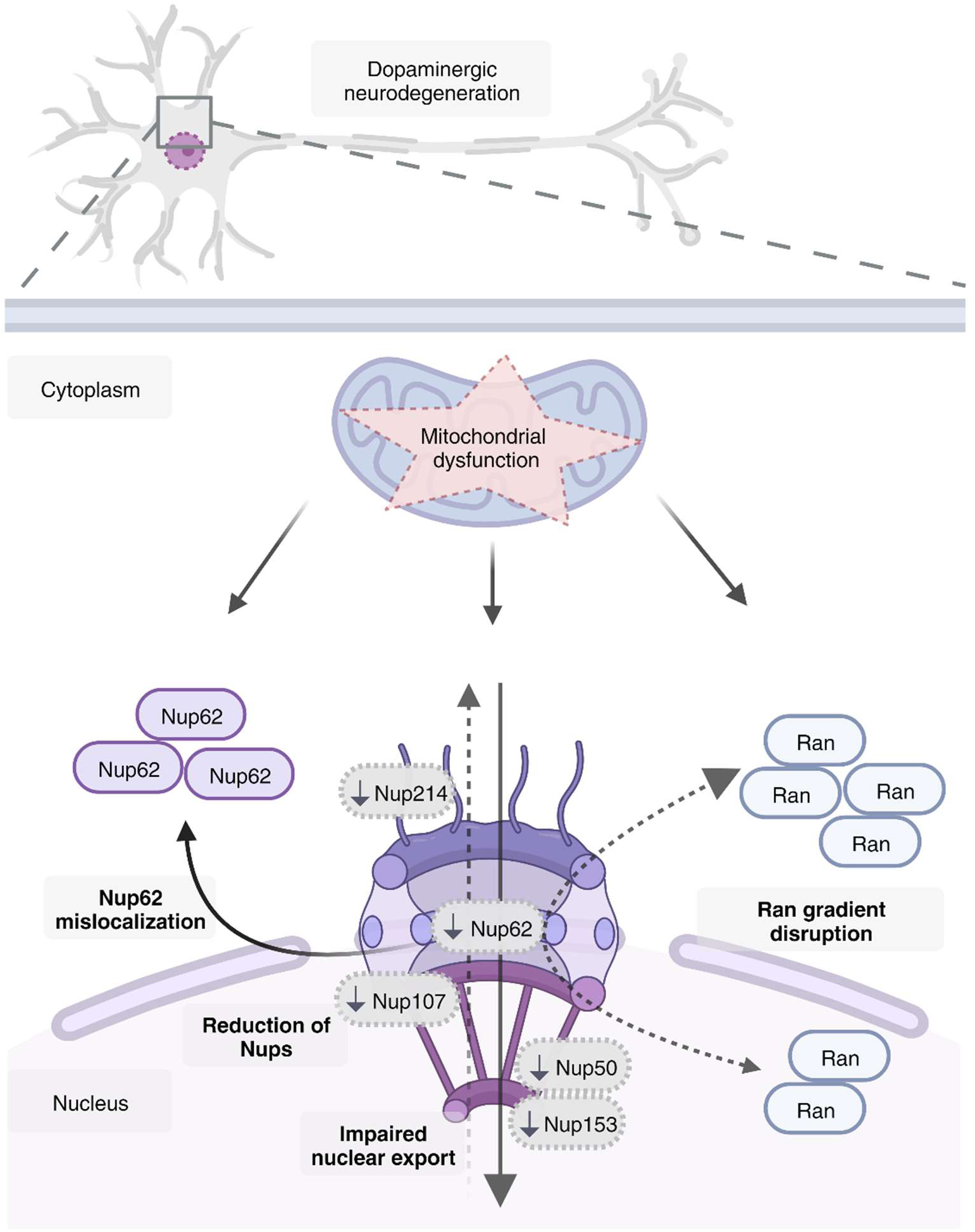

## Introduction

Parkinson’s disease (PD) is a progressive neurodegenerative disorder characterized by a loss of dopaminergic (DAergic) neurons within the substantia nigra (SN), leading to a depletion of striatal DA levels (1). Most PD cases are idiopathic, and a combination of genetic and environmental risk factors contributes to its multifactorial etiology (2). Mitochondrial dysfunction is a common pathogenic mechanism in the development of sporadic as well as familial PD (3). Exposure to neurotoxic pesticides, like the mitochondrial complex 1 inhibitor rotenone, has strongly been linked to PD-associated DAergic neuron loss (4–6). Rotenone-induced mitochondrial dysfunction leads to ATP depletion, generation of reactive oxygen species, and oxidative stress (7). While the role of mitochondrial dysfunction and oxidative stress in DAergic neurodegeneration is well established, the precise mechanisms underlying this pathophysiology remain poorly understood.

Neurons are long-lived cells with highly polarized cellular organization, which makes them particularly sensitive to disruption of the nucleocytoplasmic transport (NCT) of signaling proteins and transcription factors (8, 9). Since little to no turnover of the nuclear pore complex (NPC) occurs in post-mitotic cells, it is likely to deteriorate with accumulating oxidative damage in normal and pathological aging (10, 11). NPC damage and NCT defects have been observed in several neurodegenerative diseases (8, 12). NPCs are aqueous channels in the nuclear envelope that are formed by multiple copies of about 30 nucleoporins (Nups) intricately assembled as stable subcomplexes. They have a membrane-embedded central structure consisting of the scaffold and central channel that is skirted by the outer rings and peripheral filaments that extend into the nuclear interior and the cytoplasm (13, 14). Nups are divided into functional classes according to the subcomplex they form. Approximately one-third of the Nups contain unstructured domains rich in tandem phenylalanine and glycine repeats (FG-Nups) (15). While small molecules can pass freely through NPCs, these channels mediate the exchange of macromolecular proteins and RNA across the nucleoplasm and cytoplasm via active transport (16). This active transport is also mediated by transport receptors that recognize protein cargo containing either a nuclear localization signal (NLS) or nuclear export signal (NES). Transport receptors carrying protein cargo interact with Nups to traverse the NPC central channel (16–18). The NCT of most proteins depends on the GTP-binding nuclear protein Ran. The nuclear import complex is disassembled in the presence of RanGTP inside the nucleus, releasing the protein cargo. The nuclear export complex is formed only in the presence of nuclear RanGTP, and it is disassembled in the cytoplasm when RanGTP is hydrolyzed to RanGDP (19, 20). Ran is primarily found in the nucleus, and the nuclear-to-cytoplasmic Ran gradient is crucial for the directionality of active transport across the NPC (21). The NPC also serves several transport-independent functions including the regulation of genome organization and gene expression (22).

Specific macromolecules including transcription factors and kinases are mislocalized in DAergic neurons in PD, which implicates NCT impairment in disease pathogenesis (8, 23–26). It has been shown that α-synuclein, the neuronal protein linked to PD both genetically and neuropathologically, is involved in NCT through its interaction with Ran (27). Mutations in the most frequent PD-linked genes, *LRRK2* and *SNCA*, disrupt the nuclear lamina and nuclear envelope architecture (27–29). Moreover, it has been reported that haploinsufficiency of Nup358/Ran-binding protein 2 in mice treated with the Parkinsonian neurotoxin MPTP induced a stronger disease phenotype and slower recovery (30). Studies have shown that oxidative stress alters Nups and induces NPC damage (31, 32). Therefore, tissues with high oxidative stress levels, like the DAergic neurons, may be particularly susceptible to NPC damage (9). However, the involvement of specific Nups and NCT machinery in mitochondrial dysfunction-related DAergic neurodegeneration in PD pathogenesis has not been investigated.

In this study, we examined the distribution of specific Nups in human PD and mitochondrially impaired transgenic mouse brains, as well as in several DAergic neuronal models of mitochondrial dysfunction. We show that Nup expression is reduced in human PD neuronal nuclei, and that mitochondrial dysfunction induces downregulation of Nups in DAergic neurons in vitro and in vivo. In addition to its nuclear depletion, we found cytoplasmic mislocalization and accumulation of one FG-Nup, Nup62, in mitochondrial dysfunction models and PD brains. Furthermore, we report that mitochondrial impairment induces disruption of the Ran gradient and classical nuclear export in DAergic neurons. Ran is also depleted in the nuclei of PD patient neurons. Our in vitro and in vivo data provide direct evidence for a key role of Nup pathology in mitochondrial dysfunction-mediated DAergic neurodegeneration. Ultimately, these findings link NPC and NCT dysregulation to neuronal cell death in the pathogenesis of PD.

## Results

### Mitochondrial dysfunction triggers downregulation of specific Nups in DAergic neural cells

Nups have been demonstrated to vary in their turnover and exchange rate, structural domains, and functions depending on their location within the NPC (33). To examine any potential changes in the expression and distribution of Nups in DAergic neural cells in response to mitochondrial dysfunction, we first used a rotenone-based DAergic cell model. We previously demonstrated that exposing N27 DAergic neural cells to 1 µM rotenone, a potent mitochondrial complex I inhibitor, for up to 6 h leads to bioenergetic failure, ATP depletion, and reduced respiratory capacity without significant cell death (34, 35). In this study, we exposed N27 cells to 1 µM rotenone for 3 and 6 h, extracted nuclear and cytoplasmic fractions (Supp. Fig. 1A), and measured specific Nup levels by Western blot. To systematically examine the nuclear pore, we selected specific Nups that belong to different subcomplexes within the NPC: Nup107 from the outer ring, Nup62 from the central channel, and NUPs 50 and 153 from the nuclear basket (Fig. 1A). Our Western blot analysis reveals that rotenone exposure time-dependently downregulated Nup153, Nup107, Nup62 and Nup50 in the nuclear fraction (Fig. 1B, C). In addition, we did not observe significant mislocalization of these Nups to the cytoplasmic fraction in rotenone-treated cells (Supp. Fig. 1B, C). We found that the reduction in the protein levels of these Nups in rotenone-treated N27 cells does not correspond to a decrease in their mRNA levels (Supp. Fig. 1D).

**Fig. 1.**
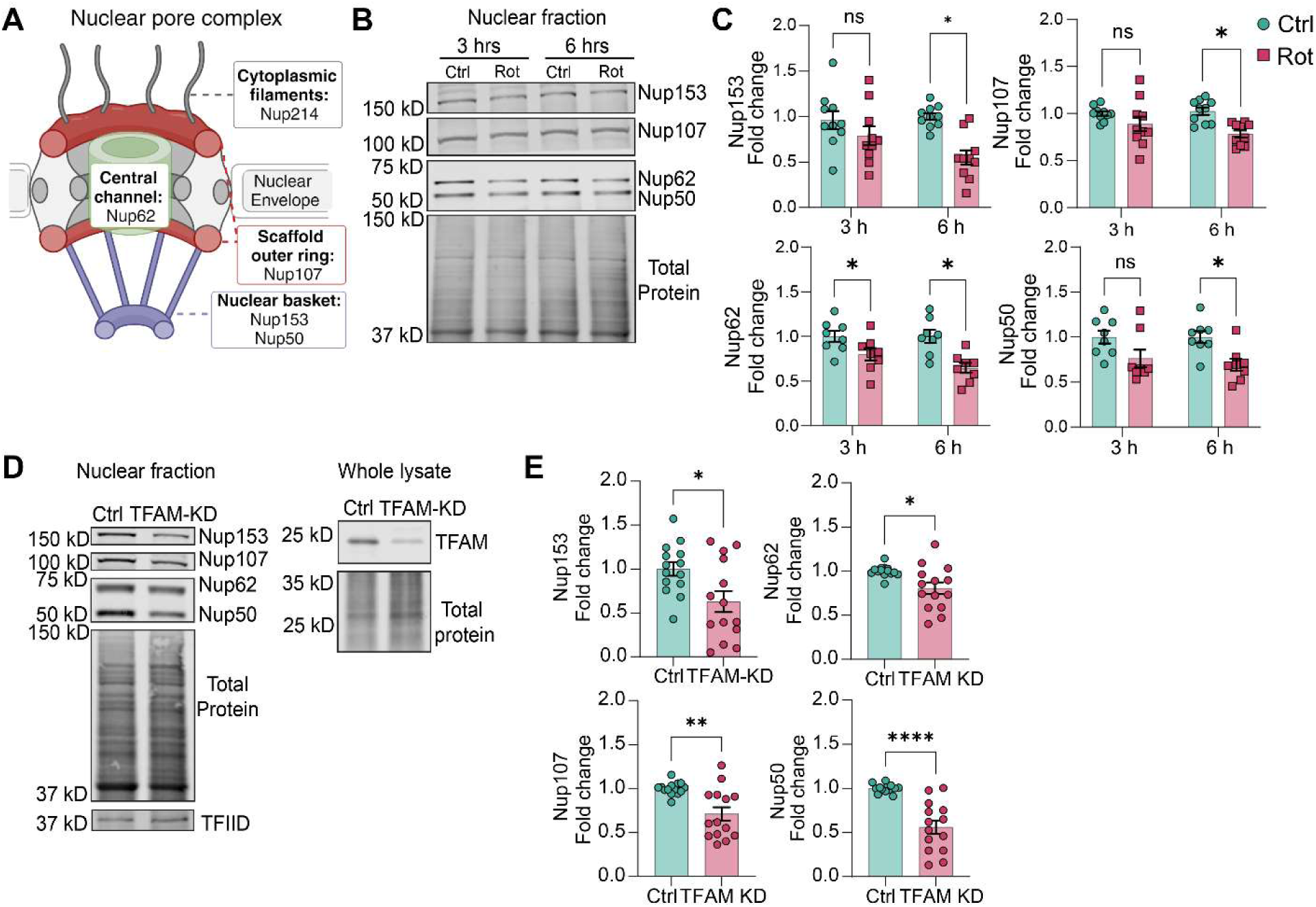
Mitochondrial dysfunction induces downregulation of Nups in N27 dopaminergic neural cells. (A) Illustration of the NPC showing Nups 214, 62, 107, 153 and 50 in their subcomplexes. Created with BioRender.com. (B) Representative Western blot images of Nups 153, 107, 62, and 50 in nuclear fraction of N27 DAergic neural cells exposed to 1µM rotenone (Rot) for 3 and 6 h and their (C) quantification from at least 4 independent experiments, with 2 technical repeats each. Multiple unpaired t-tests with Holm-Sidak method to correct for multiple comparisons. (D) RepresentativeWwestern blot images of Nups 153, 107, 62, and 50 in the nuclear fraction of CRISPR control (ctrl) and TFAM knockdown (TFAM-KD) N27 cells and TFAM levels in whole lysates of these cells. (E) Quantification of Nups in nuclear fraction of TFAM-KD N27 cells from 7 independent experiments with 2 technical repeats each. Unpaired t-tests with Welch’s correction. All western blots were normalized to total protein and Nup levels are shown as fold change relative to control (mean ± SEM). *p<0.05; **p<0.01; ****p<0.0001; ns, not significant.

To validate our finding of Nup reduction upon rotenone-induced mitochondrial stress, we used a CRISPR/Cas9-based mitochondrial transcription factor A knockdown (TFAM-KD) N27 cell culture model of mitochondrial dysfunction. TFAM is essential for mitochondrial DNA transcription and maintenance of mitochondrial DNA copy number (36). TFAM KD in N27 cells impairs mitochondrial function, alters mitochondrial morphology, and depletes cellular ATP levels (34, 35). Consistent with the findings in rotenone-exposed N27 cells, Nup153, Nup107, Nup62 and Nup50 were significantly reduced in the nuclear fraction of TFAM-KD N27 cells compared to CRISPR control cells (Fig. 1D, E). Next, we performed super-resolution structured illumination (SR-SIM) immunofluorescence analysis for a clear resolution of the distribution of Nups. Specifically, we immunostained Nup153, 107, 62 and 50, as well as the cytoplasmic filament Nup214, in N27 cells exposed to 1 µM rotenone for 6 h. Cell Profiler pipelines were used to quantify the integrated intensity of fluorescence signal from the SR-SIM images. We found significantly reduced signal intensities of Nup153, Nup107, Nup62, Nup50 and Nup214 in rotenone-treated cells (Fig. 2A-J).

**Fig. 2.**
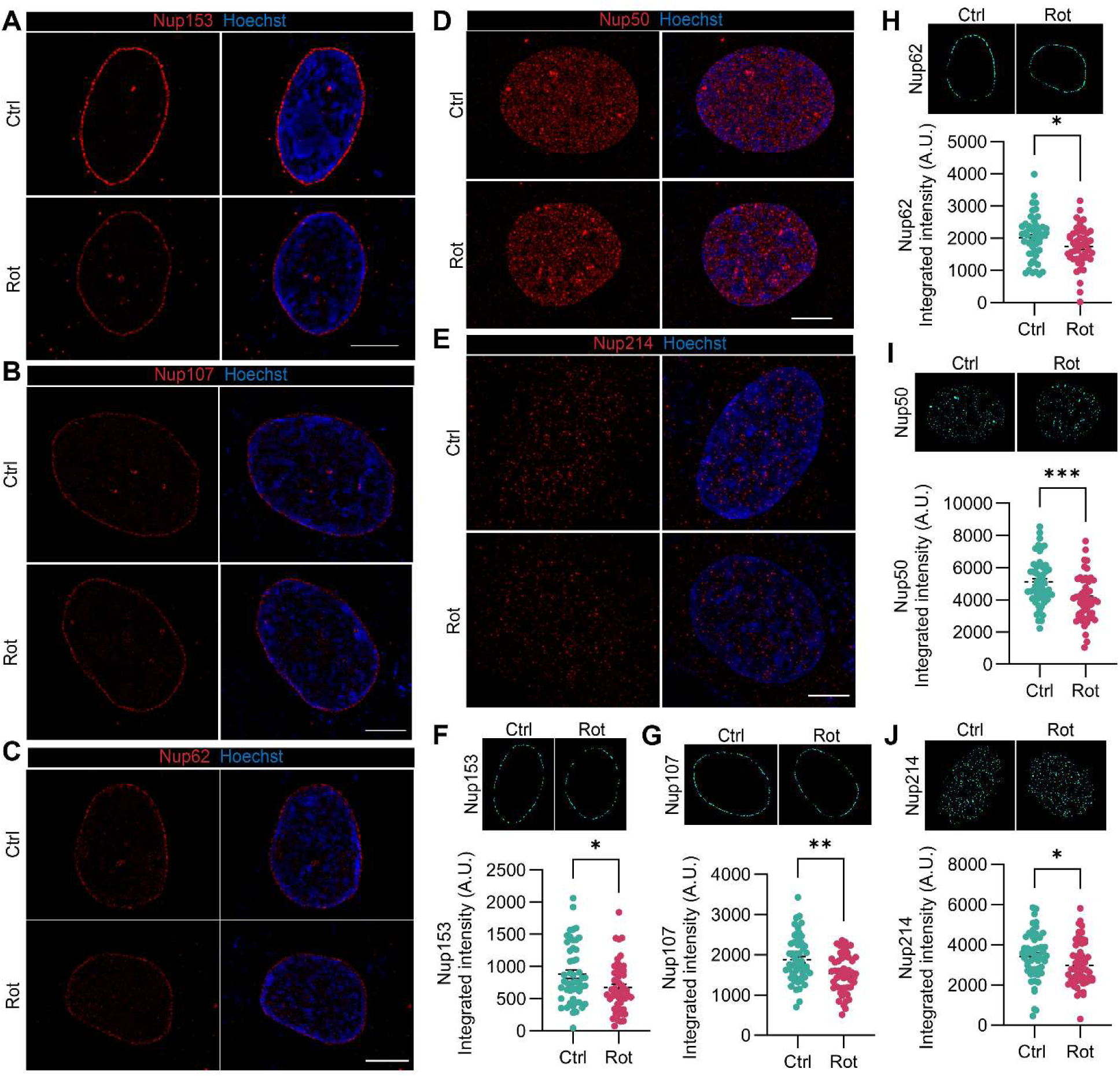
Super resolution structured illumination microscopy analysis reveals reduction in nuclear levels of Nups in N27 dopaminergic neural cells under mitochondrial stress. Representative single z slice super resolution structured illumination (SR-SIM) images of (A) Nup153, (B) Nup107 and (C) Nup62 and maximum intensity projections from SR-SIM imaging of (D) Nup50 and (E) Nup214 from immunofluorescence analysis of control and rotenone-exposed N27 DAergic neural cells. Scale bars, 5 µm. Quantification of Nup integrated signal intensity and CellProfiler subplots representing the fluorescence area that was quantified for (F) Nup153, (G) Nup107, (H) Nup62, (I) Nup50 and (J) Nup214. ≥ 50 cells were imaged for each Nup from at least 4 independent experiments. All data are presented as mean ± SEM and each data point represents one nucleus. Unpaired t-tests with Welch’s correction for unequal variances. *p<0.05; **p<0.01; ***p<0.001.

To further confirm the role of oxidative stress in Nup reduction in N27 cells, we checked nuclear levels of Nups 153, 107, 62 and 50 in N27 cells exposed to hydrogen peroxide. We observed significant reductions in Nup153, Nup62 and Nup50, but not in Nup107 in these cells (Supp. Fig. 2A, B). Overall, the nuclear basket Nups, Nup50 and Nup153, showed the greatest fold decrease relative to control cells for both rotenone-exposed and TFAM-KD N27 cell culture models. To compare the NPC morphology at high resolution between control and rotenone-exposed N27 cells, we examined their nuclear membranes using transmission electron microscopy. Interestingly, the nuclear membrane is not notably destroyed, and the NPC appears unchanged in rotenone-exposed cells compared to control cells (Supp. Fig. 3). Collectively, these findings indicate that mitochondrial dysfunction mechanistically leads to the reduction of specific Nups in DAergic neural cells in vitro, without causing an overall collapse in NPC structure.

### Nup153 is reduced in the nuclei of hiPSC-derived midbrain DAergic neurons upon mitochondrial stress

To validate our observation of rotenone-induced NPC alterations in a physiologically relevant human in vitro model, we used human induced pluripotent stem cells (hiPSCs) in which Nup153 is endogenously tagged with mEGFP at the N-terminus (37). The tag allowed us to clearly observe Nup153 alterations without relying on immunostaining. The hiPSCs were differentiated into TH^+^ and β3-Tubulin^+^ post-mitotic DAergic neurons (Supp. Fig. 4). Previously, Neeley et al. showed that exposing hiPSC-derived midbrain DAergic neurons to 1 µM rotenone for 24 h leads to significant neurite loss (38), and at 100 µM rotenone treatment, Shaltouki et al. reported overt swelling of mitochondria and loss of mitochondrial matrix density (39). At 35-40 days post differentiation, we exposed the Nup153-EGFP tagged DAergic neurons to 100 µM rotenone for 24 h. We used the higher acute rotenone dose since we treated the cells in complete differentiation media containing antioxidant compounds, which make DAergic neurons more resistant to toxicants. Upon rotenone exposure, the DAergic neurons exhibited notable neurite loss (Fig. 3A). Super-resolution imaging revealed a pronounced reduction in the Nup153-EGFP signal, no mislocalization or redistribution of Nup153 occurred (Fig. 3B). Quantification of the EGFP signal within TH^+^ neurons in fluorescence microscopy images shows a significant reduction in Nup153 upon rotenone exposure (Fig. 3C, D). Collectively, these data show that mitochondrial dysfunction significantly reduces Nup153 levels and thus alters the NPC in a human stem cell-derived model.

**Fig. 3.**
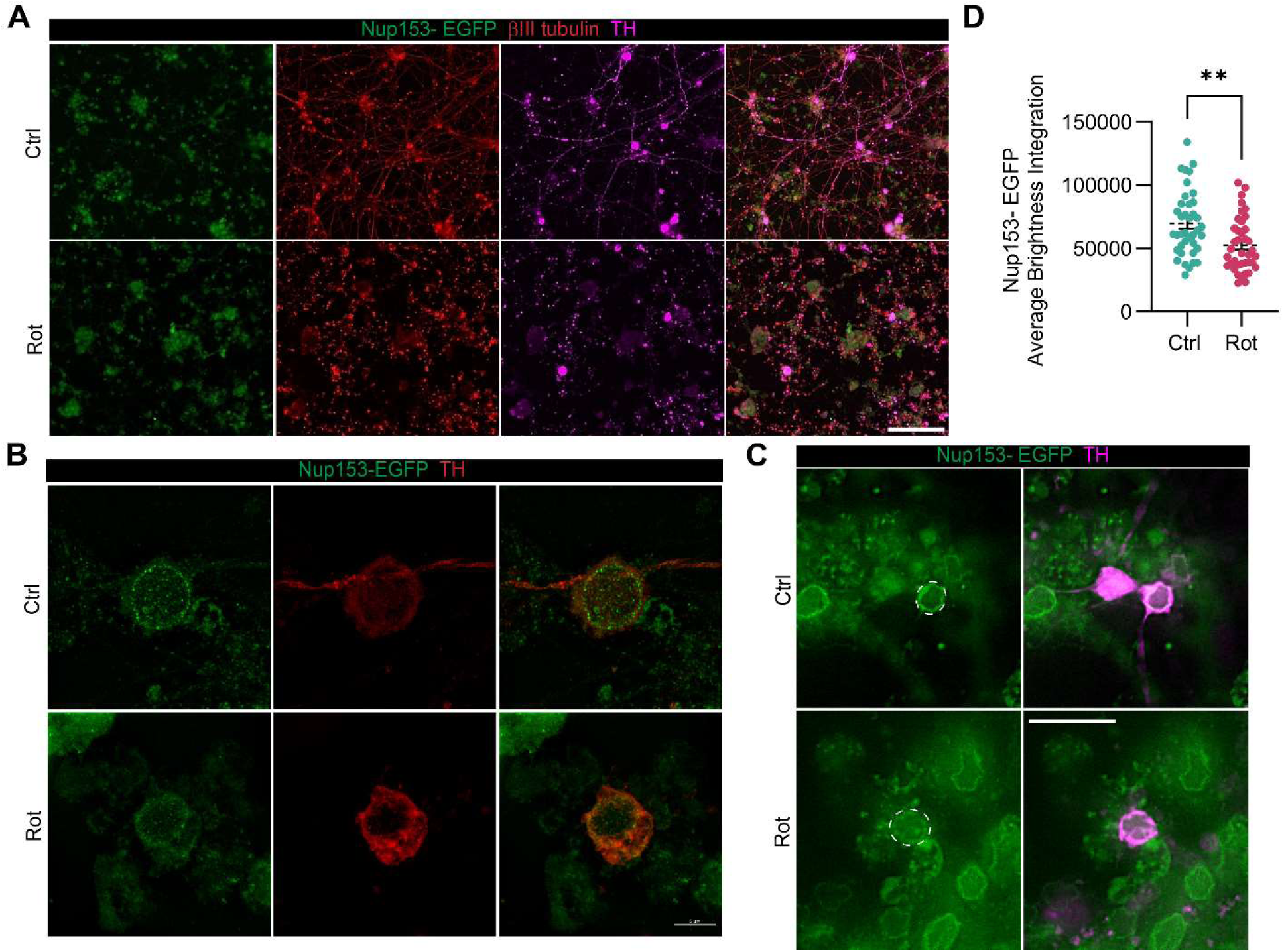
Nup153 is reduced in hiPSC-derived midbrain dopaminergic neurons upon exposure to rotenone. (A) Representative images of hiPSC derived DAergic neurons after 30-35 days of differentiation expressing Nup153-EGFP and immunostained with the neuronal marker βIII tubulin, as well as the DAergic neuron-specific marker TH. Scale bar, 100 µm. (B) Representative maximum intensity projection SR-SIM images of Nup153-EGFP in DAergic neurons labeled with TH. Scale bar, 5 µm. (C) Representative fluorescence microscopy images and (D) quantification of Nup153-EGFP integrated fluorescence intensity from TH positive neurons in these images. Scale bar, 20 µm. Data from 4 independent experiments are presented as mean ± SEM and each data point represents an image. Unpaired t-test, **p<0.01.

### Specific Nups are altered in the nigral DAergic neurons of MitoPark transgenic mice

To investigate the changes in Nups in an *in vivo* model of mitochondrial dysfunction, we used the MitoPark mouse model. This model was created by selective TFAM knockdown in DAergic neurons in the nigrostriatal pathway using the DA transporter promoter (36). MitoPark mice have been extensively characterized by our lab and other groups and they mimic key pathological features of PD, including motor deficits that begin at 12 weeks of age and become progressively worse (35), and PD-associated non-motor symptoms (40). SN sections from MitoPark mice over 20 weeks old and age- and sex-matched littermate control mice were double immunostained for Nups and the DAergic neuron marker, TH. Confocal imaging shows reduced signal of Nup153 (Fig. 4A), Nup50 (Fig. 4B), and Nup107 (Fig. 4C) in the surviving TH neurons in the SN of MitoPark mice compared to littermate controls. We also observed irregular nuclear membrane morphology and deep nuclear invaginations in the nuclei of DAergic neurons in MitoPark mice when stained with Nup153 (Fig. 4A). Upon quantification of the nuclear signal intensity of these Nups from fluorescence microscopy images (Supp. Fig. 5A-C), we observed a significant reduction in Nup153 (Fig. 4D), Nup50 (Fig. 4E), and Nup107 (Fig. 4F). Next, we checked the changes in the general distribution of NPC proteins in the nuclei of MitoPark mice by staining SN sections with the mAb414 antibody that recognizes FG-repeat-containing Nups including Nups 358, 214, 153 and 62. Confocal imaging did not reveal any substantial change in mAb414 signal distribution (Fig. 4G); instead, we observed a significant increase in mAb414 signal intensity in fluorescence microscopy images of DAergic neurons in MitoPark mice compared to littermate controls (Fig. 4H, Supp. Fig. 5D). This suggests that without an overall reduction in NPCs, or change in NPC distribution, the relative abundance of specific Nups in the nuclear membrane of DAergic neurons gets reduced upon mitochondrial stress in vivo.

**Fig. 4.**
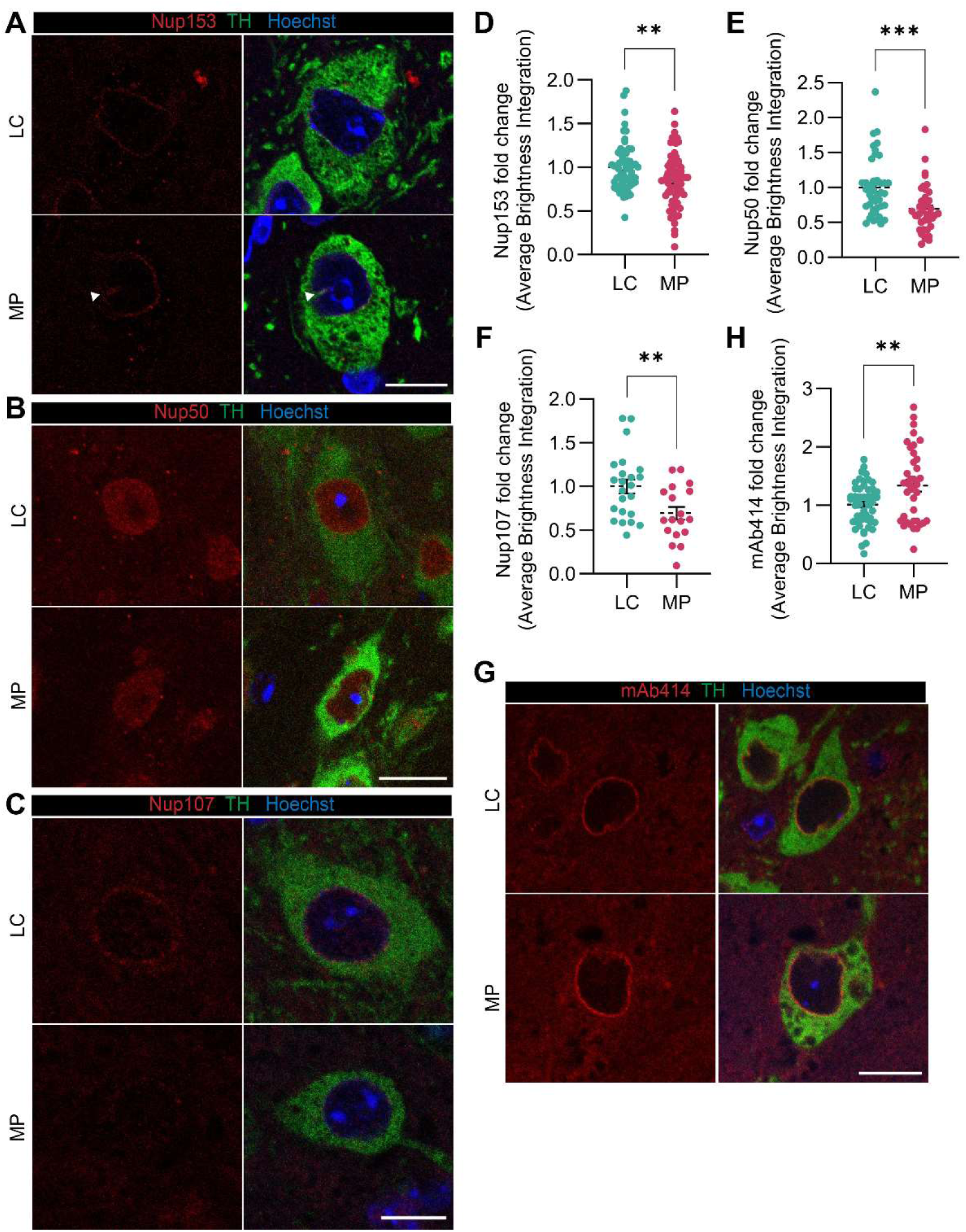
Specific Nups are reduced in substantia nigra of MitoPark transgenic mice. Representative confocal images of SN sections of MitoPark (MP) and littermate control (LC) mice immunostained for TH and (A) Nup153, (B) Nup50, and (C) Nup107. Arrowhead in (A) shows irregular contours and invagination of the nuclear membrane. Quantification of (D) Nup153, (E) Nup50, and (F) Nup107 signal. (G) Representative confocal images and (H) quantification of the NPC stained by mAb414 antibody. Integrated intensity of the Nup signals was quantified from fluorescence microscopy images (Supp. Fig. 5) and is shown as fold change relative to LC with 5-6 mice per group. All data are presented as mean ± SEM and each data point represents one image. Unpaired t-tests with Welch’s correction for unequal variances, **p<0.01; ***p<0.001. Scale bars, 10 µm.

For the NPC central channel component FG-Nup, Nup62, we noted an altered staining pattern in confocal images of the SN in MitoPark mice. Like Nup153 staining, Nup62 staining showed prominent nuclear invaginations in the nuclei of DAergic neurons. Additionally, a notable amount of Nup62 mislocalized from the neuronal nuclear membrane and accumulated in the cytoplasm (Fig. 5A, Supp. Fig. 6A). Quantification of Nup62 immunoreactivity in fluorescence microscopy images showed a significant reduction in Nup62 signal intensity within the nuclei of TH neurons in MitoPark mice, while the signal intensity in the cytoplasm was significantly increased (Fig. 5B). Interestingly, Nup62 mislocalization was only seen in nigral DAergic neurons and not in the hippocampal region (Fig. 5C). We checked if the cytoplasmic Nup62 aggregates are ubiquitinated since the ubiquitin-proteasome system is one of the pathways that is impacted in PD pathology (41), especially given the evidence for crosstalk between mitochondrial impairment and proteasome dysfunction (42, 43). However, we did not observe colocalization of Nup62 and ubiquitin signal in the double immunostained SN sections of MitoPark mice (Supp. Fig. 6B).

**Fig. 5.**
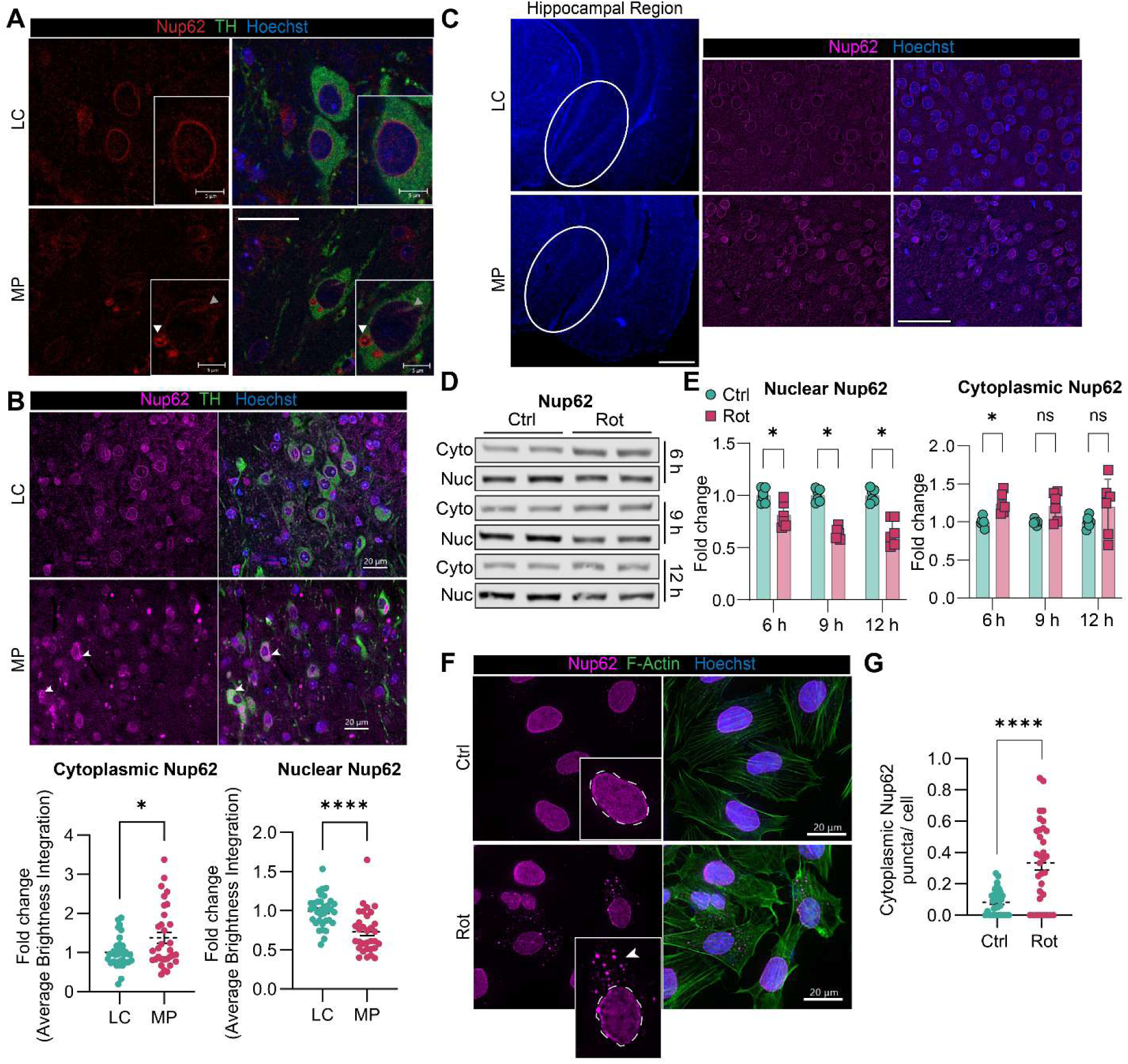
Mitochondrial dysfunction promotes cytoplasmic mislocalization and accumulation of Nup62 specifically in dopaminergic neurons. (A) Representative confocal images of SN sections of MitoPark (MP) and littermate control (LC) mice immunostained for TH and Nup62. White arrowhead shows cytoplasmic staining of Nup62, and grey arrowhead indicates nuclear membrane invagination. Scale bars, 10 µm; insets, 5 µm. (B) Representative fluorescence microscopy images and quantification of nuclear and cytoplasmic Nup62 fluorescence integrated intensity from these images. Arrows show cytoplasmic staining of Nup62 in MP SN. Scale bars, 20 µm. Quantification is shown as fold change relative to LC for 6 mice per group. Data are presented as mean ± SEM and each data point represents one image. Unpaired t-tests with Welch’s correction for unequal variances. (C) Nup62 immunostaining in the hippocampal region of MitoPark mice. Scale bar in zoomed out image of hippocampal region, 200 µm; zoomed images, 20 µm. (D) Western blot images and (E) quantification of Nup62 in cytoplasmic and nuclear fractions of N27 DAergic neural cells exposed to low-dose (100 nM) rotenone for 6, 9 and 12 h. Nup62 bands were normalized to total protein (Supp. Fig. 7B). Data shown as fold change relative to control from 3 independent experiments with 2 technical replicates each (mean ± SEM). Multiple unpaired t-tests with Holm-Sidak method to correct for multiple comparisons. (F) Representative immunofluorescence images of Nup62 in N27 cells exposed to low-dose rotenone for 12 h. Arrowhead shows cytoplasmic Nup62. Scale bars, 20 µm. (G) Quantification of cytoplasmic Nup62 puncta per cell. Data from 3 independent experiments are presented as mean ± SEM and each data point represents one image. Unpaired t-test with Welch’s correction. *p<0.05; ****p<0.0001; ns, not significant.

### Low-dose rotenone exposure induces cytoplasmic mislocalization of Nup62 in DAergic neural cells

Based on our in vivo observation of Nup62 accumulation in the cytoplasm due to chronic and prolonged mitochondrial stress, we checked if a low dose and less acute rotenone treatment will lead to Nup62 mislocalization in N27 DAergic neural cells as well. For this purpose, we treated N27 cells with 100 nM rotenone for 6, 9 and 12 h. Our Western blot of Nup62 levels in the nuclear and cytoplasmic fractions of these cells reveals a significant, time-dependent reduction in nuclear Nup62, and a significant increase in cytoplasmic Nup62 after 6 h of rotenone treatment (Fig. 5D, E). Immunofluorescence analysis of Nup62 in N27 cells exposed to 100 nM rotenone for 12 h also shows a significant three-fold increase in Nup62 cytoplasmic puncta per cell (Fig. 5F, G). Previous studies report that Nup62 contributes to pathologic aggregates and localizes to stress granules during neurodegeneration (44–46). To check whether the cytoplasmic Nup62 puncta in low-dose rotenone-exposed cells are stress granule assemblies, we stained for Nup62 and the stress granule marker G3BP1. However, G3BP1 and cytoplasmic Nup62 puncta staining did not overlap, suggesting that the cytoplasmic Nup62 does not localize to G3BP1-containing stress granules in these cells (Supp. Fig. 7A). We also examined Nup153, Nup107 and Nup50 levels in N27 cells exposed to 100 nM rotenone. Nuclear levels of these Nups were significantly reduced in a time-dependent manner (Supp. Fig. 7B, C). Nup50 and Nup153 levels were not significantly altered in the cytoplasmic fraction, but cytoplasmic Nup107 significantly increased after 12 h of exposure to 100 nM rotenone (Supp. Fig. 7D). However, immunofluorescence analysis of Nup107 in N27 cells exposed to 100 nM rotenone for 12 h did not show a cytoplasmic signal (Supp. Fig. 7E). Collectively, these data show that Nups time-dependently decrease in the nuclei of N27 cells upon exposure to low-dose rotenone, and Nup62 mislocalizes to the cytoplasm and forms aggregates.

### Nup153 and Nup62 are altered in postmortem PD patient brain tissues

To identify whether Nup pathology is a feature of PD pathogenesis in human patients, we compared Nup153 and Nup62 levels and localization in human postmortem PD brains and age-matched healthy control brains. We first checked Nup153 and Nup62 levels in whole lysates from fresh frozen SN tissues (Fig. 6A). Our findings revealed a trend towards a decrease in Nup153 levels, while Nup62 significantly increased 0.5-fold in PD SN compared to control (Fig. 6B). Next, we immunostained fresh frozen SN sections for Nup153 and 62. Consistent with the Western blot analysis, Nup153 decreased in the TH neurons of PD sections (Fig. 6C, D). Interestingly, Nup62 was depleted in the nuclei but formed extranuclear aggregates (Fig. 6E, F), which likely contribute to the increase in Western blot signal from whole tissue lysates. Collectively, these data show that Nup pathology is present in DAergic neurons from PD patient tissue and add clinical relevance to our observations in mitochondrial dysfunction induced in vitro and in vivo PD models.

**Fig. 6.**
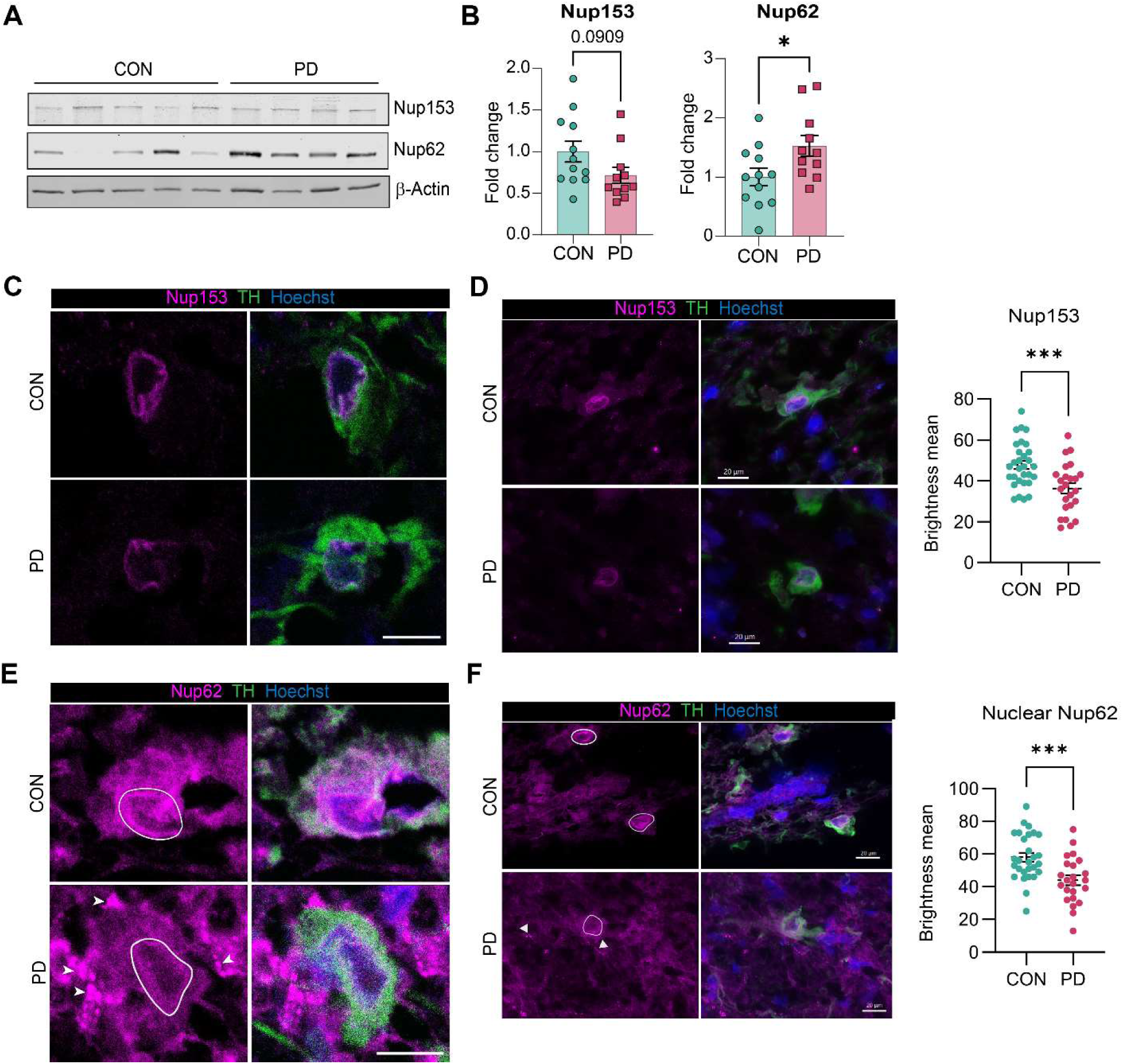
Presence of Nup pathology in SN tissue from PD patients. (A) Western blot images and (B) quantification of Nups 153 and 62 in postmortem SN tissue lysates from non-disease control (CON) and PD patients (n= 12 CON and 11 PD). Data shown as fold change relative to control (mean ± SEM). (C) Representative confocal images of SN sections from non-disease control and PD patients showing Nup153 in DAergic neurons. Scale bars, 10 µm. (D) Fluorescence microscopy images and quantification of Nup153 mean fluorescence intensity from these images. Scale bars, 20 µm (n= 3 CON and PD brains). (E) Representative confocal images of SN sections from non-disease control and PD patients showing Nup62 in DAergic neurons. Nuclear Nup62 is encircled and extranuclear Nup62 staining is indicated by arrows. Scale bars, 10 µm. (F) Fluorescence microscopy images of Nup62 with nuclear Nup62 encircled and extranuclear Nup62 signal indicated by arrows. Nuclear Nup62 mean fluorescence intensity from these images was quantified. Scale bars, 20 µm. (N= 4 CON and PD brains). Unpaired t-tests, *p<0.05; ***p<0.001; p-value shown.

### Nucleocytoplasmic distribution of Ran-GTPase is disrupted in mitochondrial dysfunction-induced PD models and in PD brains

To check whether mitochondrial stress-induced Nup alteration leads to functional impairment of the NPC, we examined the nuclear-to-cytoplasmic ratio of the Ran-GTPase protein (Ran gradient), which can be used to assay the NCT of biomolecules across the nucleus (47, 48). We immunofluorescently labeled Ran in N27 DAergic neural cells exposed to 1 µM rotenone for 6 h and quantified the fluorescence intensities of Ran in the nucleus and cytoplasm. The nuclear-to-cytoplasmic Ran gradient significantly decreased in rotenone-exposed N27 cells compared to control cells (Fig. 7A – C, Supp. Fig. 8). To determine if the Ran gradient is also disrupted in DAergic neurons under mitochondrial stress in vivo, we immunostained MitoPark and littermate control sections for Ran and TH. We also observed a reduction in nuclear-to-cytoplasmic Ran ratio in TH neurons of MitoPark mice compared to control mice (Fig. 7D, E). Next, we checked Ran distribution in DAergic neurons in PD brains by immunolabeling control and PD SN sections. Our results showed significant depletion of Ran from the nuclei of TH neurons in PD brains compared to control, indicating a disruption in the Ran gradient across the nuclear membrane (Fig. 7F, G). These data show that mitochondrial dysfunction disrupts the Ran gradient in DAergic neurons in vitro and in vivo, and that Ran is aberrantly distributed in PD neurons.

**Fig. 7.**
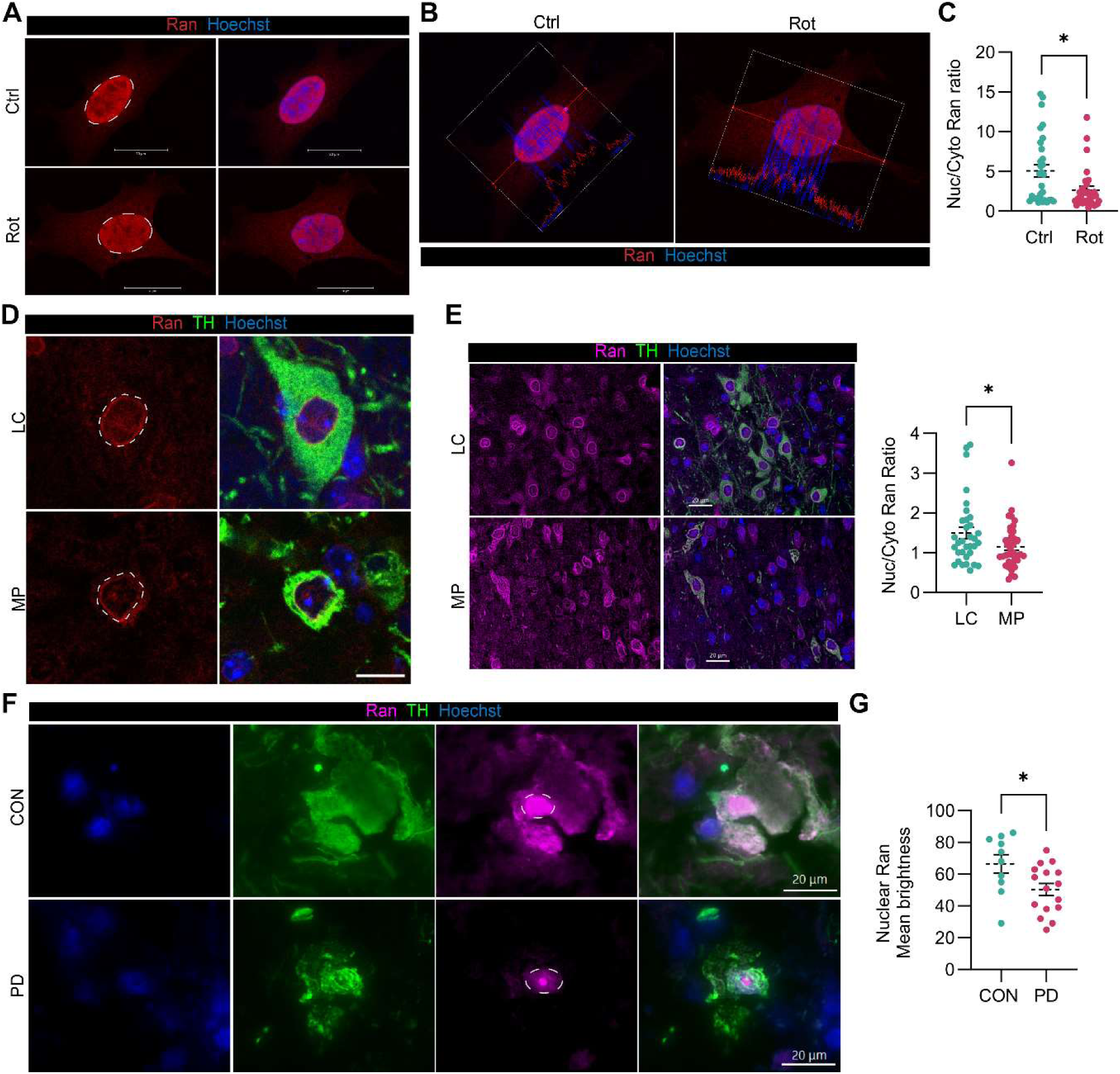
Mitochondrial stress disrupts nucleocytoplasmic transport in N27 dopaminergic neural cells. (A) Representative confocal images and (B) intensity profile of Ran distribution in control and rotenone exposed N27 DAergic neural cells. Scale bars, 20 µm. (C) Quantification of nuclear/ cytoplasmic Ran distribution in control and rotenone exposed N27 cells from fluorescence microscopy images (Supp. Fig. 8A). Data from 3 independent experiments are presented as mean ± SEM and each data point represents the Ran ratio from one image. (D) Representative confocal images of Ran distribution in nigral DAergic neurons of MitoPark (MP) and littermate (LC) control mice. Dotted outlines highlight the depletion of nuclear Ran from DAergic neuron in MP SN. Scale bars, 10 µm. (E) Representative fluorescence microscopy images and quantification of nuclear to cytoplasmic Ran gradient in LC and MP SN sections. Scale bars, 20 µm. Data shown as fold change relative to LC for 7 mice per group (mean ± SEM) and each data point represents one image. (F) Representative immunofluorescence images of Ran distribution and (G) quantification of nuclear Ran intensity in DAergic neurons from SN sections of non-disease control (CON) and PD patients (n= 3 CON and PD brains). Scale bars, 20 µm. Unpaired t-tests with Welch’s correction, *p<0.05.

### Mitochondrial stress impairs nuclear export in DAergic neural cells

To further investigate the effect of mitochondrial dysfunction on the NCT of proteins, we used a dual fluorescent reporter system. This reporter is in a lentiviral mammalian expression plasmid and comprises GFP and RFP fused to nuclear export and localization signals (GFP-NES and RFP-NLS), respectively, with an internal ribosome entry site between them (49). In healthy neurons, we expect GFP-NES to be excluded from the nucleus and RFP-NLS to be localized in the nucleus. Conversely, in the case of impaired NCT, GFP-NES and RFP-NLS would be homogenously distributed in the cell (Fig. 8A). We expressed the fluorescent reporters in N27 DAergic neural cells and then exposed them to 1 µM rotenone for 6 h. We measured the GFP and RFP signal intensities and quantified the nuclear-to-cytoplasmic ratios of these proteins. In control cells RFP-NLS is localized in the nucleus, and GFP-NES remains in the cytoplasm. In rotenone-exposed cells, although the nuclear-to-cytoplasmic RFP-NLS ratio did not significantly change, we observed a modest but significant increase in the nuclear-to-cytoplasmic GFP-NES ratio (Fig. 8B, C). Collectively, these results suggest that rotenone treatment impairs nuclear export but not nuclear import, resulting in the mislocalization of GFP-NES protein within the cell (Fig. 8D).

**Fig. 8.**
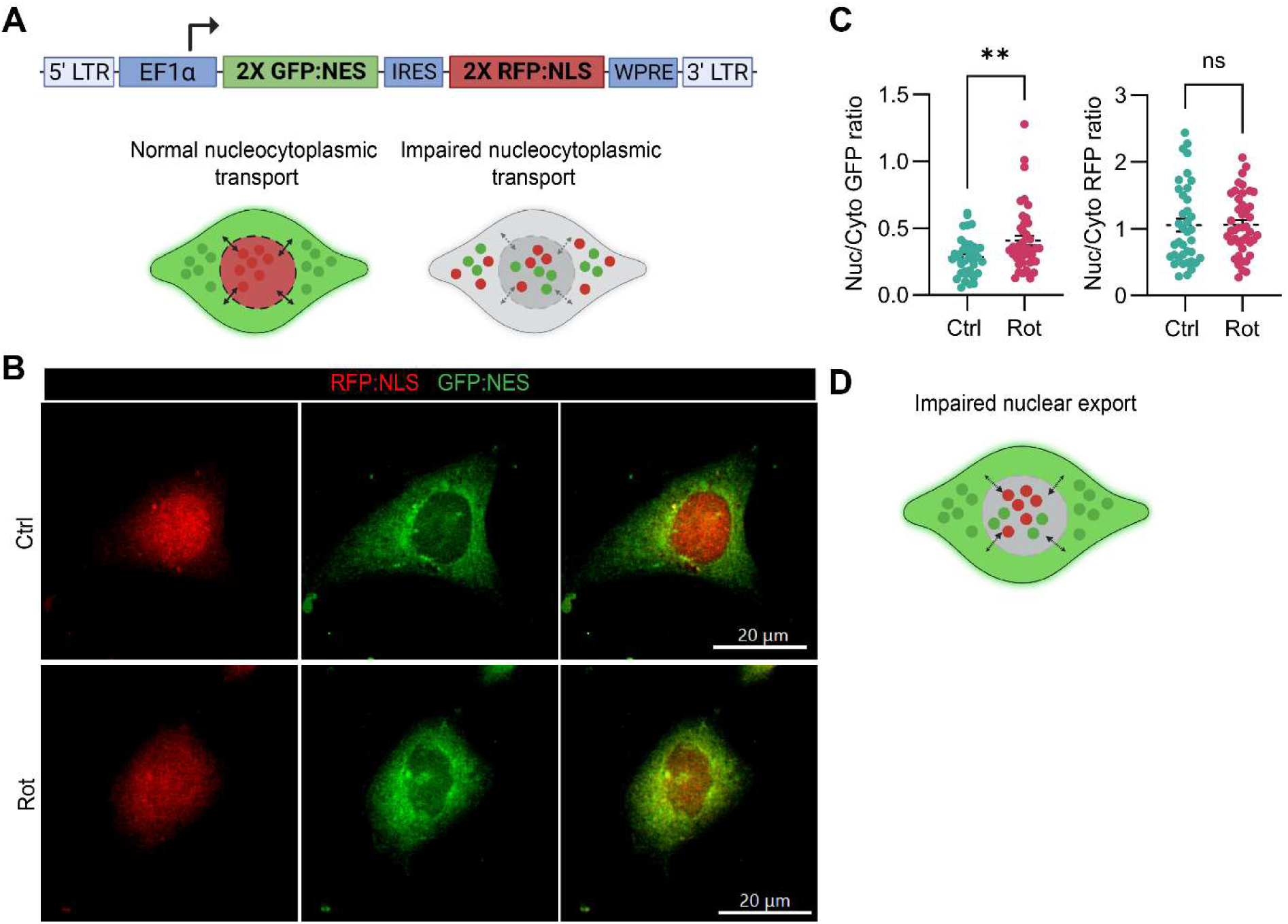
Ran gradient is disrupted in SN tissues from MitoPark mice and PD patients. (A) Illustration of the fluorescent reporter construct used to study nucleocytoplasmic transport and relative distributions of the reporter proteins in healthy neurons and in neurons with impaired nucleocytoplasmic transport. (B) Representative images of control and rotenone-exposed N27 cells transduced with the construct and (C) quantification of the nuclear/ cytoplasmic ratio of the GFP-NES and RFP-NLS reporter proteins. Scale bars, 20μm. (D) Illustration of disrupted nuclear export in N27 cells upon rotenone exposure. Data from 3 independent experiments are presented as mean ± SEM and each data point represents the reporter ratio from one image. Unpaired t-test with Welch’s correction, **p<0.01; ns, not significant. Illustrations created with BioRender.com.

### Co-treatment with the mitochondrially targeted antioxidant Mito-apocynin restores levels of Nup153 and 107 in DAergic neural cells exposed to rotenone

To determine if suppression of oxidative stress averts nuclear pore alteration in rotenone-exposed N27 DAergic neural cells, we used the mitochondrially targeted antioxidant, Mito-apocynin. We have previously shown that treating mitochondrially impaired N27 DAergic neural cells with 10 – 30 µM Mito-apocynin improved mitochondrial structural integrity, increased the oxygen consumption rate and ATP levels, and decreased mitochondrial oxidants (35). In this study, we co-exposed N27 DAergic neural cells to 1 µM rotenone and 30 µM Mito-apocynin. Rotenone treatment alone notably shortened the mitochondria, making them appear more circular, but Mito-apocynin co-treatment improved mitochondrial morphology, rendering them longer and more filamentous like control cells (Supp. Fig. 9). Further, SR-SIM immunofluorescence analysis revealed that co-exposing N27 cells to both rotenone and Mito-apocynin did not downregulate Nup153 (Fig. 9A, B) and Nup107 (Fig. 9C, D) unlike rotenone exposure alone. These results indicate that suppression of mitochondrial dysfunction by antioxidant supplementation may be a viable approach for restoring Nup levels and NPC function in DAergic neurons.

**Fig. 9.**
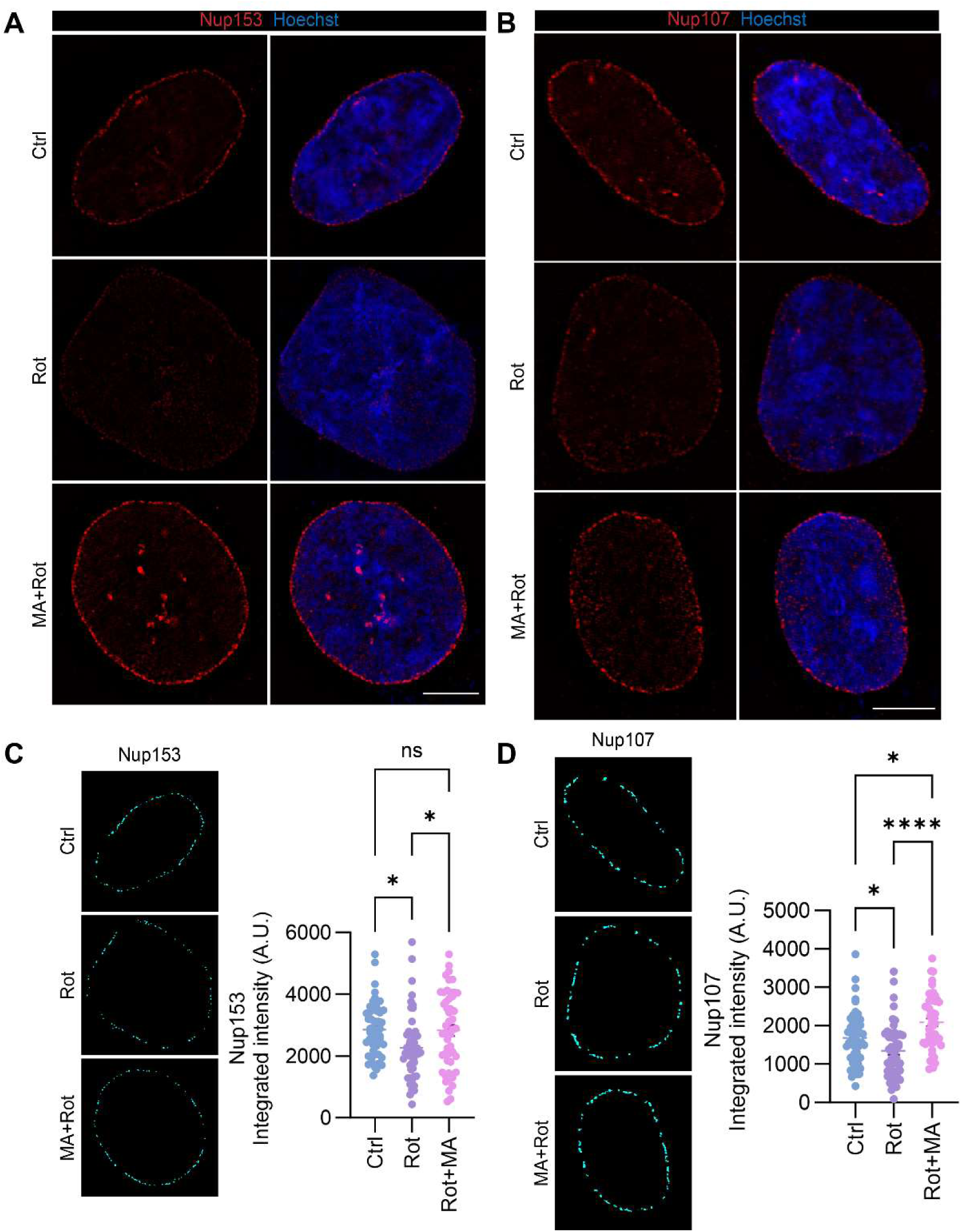
Mito-apocynin restores Nup levels in rotenone-exposed N27 dopaminergic neural cells. Representative single z slice SR-SIM images of (A) Nup153 and (B) Nup107 and from immunofluorescence analysis of control (ctrl), rotenone (rot) alone, and Mito-apocynin (MA) and rotenone-exposed N27 DAergic neural cells. Scale bars, 5 µm. Quantification of Nup integrated signal intensity and CellProfiler subplots representing the fluorescence area that was quantified for (C) Nup153 and (D) Nup107. ≥ 48 cells were imaged for each Nup from 3 independent experiments. All data are presented as mean ± SEM and each data point represents one nucleus. Ordinary one-way ANOVA with Tukey’s multiple comparisons test, *p<0.05; ***p<0.001; ns, non-significant.

## Discussion

Disruption of the NPC and impairment of bidirectional NCT are emerging as crucial pathogenic mechanisms of several neurodegenerative diseases (12, 50, 51). NPCs are extremely long-lived in post-mitotic cells (10) and tissues with high oxidative stress levels, like the DAergic neurons, may be more susceptible to NPC damage (9). Evidence of nucleocytoplasmic mislocalization of key macromolecules (8, 31) and disruption of nuclear envelope architecture (27, 28, 52) in PD models supports the need for detailed NPC analysis. Although oxidative stress has been linked to impairment of nuclear transport apparatus in other models (53, 54), it is unknown whether mitochondrial dysfunction induces NPC defects in PD-related neurodegeneration.

In this study, we provide multiple lines of evidence indicating that DAergic neurodegeneration induced by mitochondrial impairment alters the levels and distribution of specific Nups and disrupts nuclear pore function. We demonstrate that mitochondrial impairment induces a significant reduction in Nups 214, 107, 62, 153 and 50 in N27 DAergic neural cells, without an overall collapse of the NPC or a decrease in the mRNA expression of these Nups. This aligns with Coyne et al.’s findings, where a specific subset of Nups was reduced in iPSC-derived spinal neurons from patients with *C9orf72* repeat expansion-induced amyotrophic lateral sclerosis (ALS), without a decrease in their steady-state transcripts or an overall alteration of the number and distribution of NPCs (55). These Nups localize to different subcomplexes that constitute the structure of the NPC and collectively participate in NCT, chromatin regulation, and several cell signaling pathways (18, 56). Therefore, the alteration and reduction of Nups may dysregulate multiple cellular processes in DAergic neurons in response to mitochondrial impairment. Notably, the nuclear basket component Nup153 was the most consistently and drastically reduced Nup in our N27 cell models of mitochondrial dysfunction and oxidative stress whether chemically or genetically induced. We also show evidence of Nup153 loss in hiPSC-derived DAergic neurons under mitochondrial stress as well as in the nigral DAergic neurons of PD patients. Nup153 plays a crucial role in mediating the binding of export cargo within the nucleus and transport of macromolecules across the nuclear pore (57–59). In addition to its function in protein and mRNA transport, Nup153 is also involved in regulating gene expression through chromatin remodeling enzymes (60). Leone et al. previously showed that Nup153 expression is strongly reduced in neural stem cells from Alzheimer’s Disease mouse models, which alters histone 3 epigenetic marks and affects cell plasticity and neurogenesis (61). Recent work from our group has demonstrated that mitochondrial dysfunction in DAergic neurons induces histone hyperacetylation and upregulation of genes involved in neuronal apoptosis and neuroinflammation pathways (34). Although we did not investigate the interaction between Nup153 and histone modifications in this study, but others have found evidence supporting Nup153 interaction with chromatin remodelers and active chromatin marks (62, 63). Therefore, Nup153 dysfunction may be a potential mechanism contributing to mitochondrial impairment-induced epigenetic and gene expression alterations in DAergic neurons.

We report finding reduced nuclear expression of Nup107, 153 and 50 in DAergic neurons of the MitoPark mouse model of mitochondrial impairment. Interestingly, we observed a greater decrease (about 30%) in the NPC scaffold component Nup107 compared to peripheral Nups in MitoPark mouse neurons, contrasting with our cell model findings. Scaffold Nups are known to be the most long-lived and stably incorporated components of the NPC, making them more prone to age-dependent oxidative damage (9, 64). Since the MitoPark mice used in this study were aged and under prolonged oxidative stress, we speculate that the NPC scaffold may accumulate more damage than peripheral Nups. Coyne et al. have reported that Nup107 is among the small subset of Nups that are age-dependently reduced in *C9orf72* ALS patient iPSC-derived spinal neurons (55).

We noted nuclear depletion and cytoplasmic mislocalization of the NPC central channel Nup62 in vivo, specifically in the SN of MitoPark mice that we did not observe in N27 cells acutely exposed to rotenone. This suggests that Nup62 mislocalizes from the NPC to the cytoplasm in DAergic neurons when under chronic mitochondrial stress. To further examine this possibility, we exposed N27 cells to a lower dose of rotenone for a longer time. Indeed, subacute rotenone exposure led to Nup62 cytoplasmic mislocalization and accumulation in N27 cells. Besides Nup62, Western blot analysis of the cytoplasmic fraction also revealed a significant increase in another Nup, Nup107. However, unlike Nup62, we did not observe Nup107 mislocalization in immunofluorescently labeled cells, nor in the SN of MitoPark mice. This discrepancy may be due to the use of different Nup107 antibodies in the Western blot and immunostaining experiments. The cytoplasmic mislocalization of Nup62, however, appears to be consistent and clinically relevant because we found that Nup62 also accumulates outside the nuclei in human PD patient DAergic neurons. Several neurodegenerative disease studies provide evidence for Nup62 mislocalization and cytoplasmic aggregation in pathologic assemblies along with other components of the NCT apparatus as reviewed by Cristi et al. (51). A commonality of these aggregates is the presence of pathogenic proteins such as TAR DNA-binding protein 43 (TDP-43) and Fused in Sarcoma (FUS) in ALS (65, 66), and mutant Huntingtin in Huntington’s Disease (47). Aberrant phase transitions recruit Nup62 to these pathological protein inclusions (44, 67). PD is similarly categorized as a proteinopathy due to aggregation of α-synuclein in disease pathogenesis (68). However, in this study, we did not investigate the possibility of α-synuclein and Nup62 co-aggregating in human PD brains. Importantly, our findings from mitochondrial impairment models suggest that mitochondrial stress could drive Nup62 depletion from the NPC and mislocalization to the cytoplasm. Notably, while MitoPark mice have been documented to develop cytoplasmic proteinaceous aggregates containing mitochondrial protein and membrane components, these aggregates lack α-synuclein immunoreactivity (36, 69). This suggests that Nup62 mislocalizes to cytoplasmic aggregates in the absence of α-synuclein pathology. In clinical manifestations of sporadic PD, however, both mitochondrial dysfunction and proteotoxicity are cardinal pathogenic mechanisms with probable crosstalk between them (70). Future studies should explore the mechanism of Nup62 aggregation, and identify the proteins being sequestered in these aggregates in degenerating DAergic neurons.

Using the NPC marker mAb414 antibody, we demonstrate that mitochondrial impairment alters a specific set of Nups without decreasing the overall numbers and distribution of the NPC in nigral DAergic neurons of MitoPark mice. Interestingly, we note an increase in mAb414 fluorescence intensity in the SN of MitoPark mice, despite decreased levels of mAb414-reactive Nups 214, 153, and 62 in our mitochondrial dysfunction models. To explain the increase in mAb414 signal, we checked the levels of another mAb414-reactive Nup, Nup358, under mitochondrial stress in N27 DAergic neural cells. The 0.25-fold, albeit non-significant, increase in Nup358 fluorescence signal (Supp. Fig. 10) warrants further investigation of this Nup in DAergic neurons under mitochondrial stress. It has previously been documented that oxidative stress induces disulfide bond formation between Nups, leading to a global structural change of the NPC (71, 72), but more data is needed to determine if oxidative stress increases NPC numbers. Overall, our findings suggest specific downregulation of individual Nups without a reduction in NPC numbers and distribution, consistent with previous finding (55).

Defects in NCT components have been implicated as a mechanistic driver of PD pathogenesis based on the observation by Chen et al. that mutant α-synuclein sequesters Ran and prevents it from transporting cargo across the nucleus (27). Here, we have demonstrated that Ran is depleted from the nuclei of DAergic neurons in the SN of PD patient brain sections, further implicating NCT defects in disease pathogenesis. Also, our in vitro and in vivo results showing that mitochondrial dysfunction disrupts the Ran gradient in DAergic neurons are consistent with previous studies documenting that cellular oxidative stress adversely affects Ran distribution and NCT (54, 73). Ran has been shown to be a redox-active protein, making it a primary target of oxidative stress (74). Our data indicates that mitochondrial impairment in DAergic neurons is sufficient to disrupt the localization of Ran and consequently perturb NCT. The altered nucleocytoplasmic Ran gradient may result from either NPC dysfunction or direct modification of Ran by oxidants in DAergic neurons under mitochondrial stress. However, it is important to note that Ran localization is disrupted in two crucial pathogenic mechanisms of PD, α-synuclein toxicity and mitochondrial dysfunction, suggesting that NCT defect is a key contributor to disease pathogenesis. We further examined mitochondrial dysfunction-induced NCT defects using fluorescent reporter proteins and found that, while nuclear import is unaffected, nuclear export is impaired (Fig. 7). This result differs from previous studies in other neurodegenerative disease models that express mutant TDP-43 and *C9ORF72* repeat expansion, which leads to only nuclear import defects (66, 75). Additional neurodegeneration models that express mutant ALS-linked FUS and pathological tau show an overall nuclear transport defect affecting both nuclear import and export (48, 65). Studies in non-neuronal models of oxidative stress have also reported nuclear import as well as export defects (76–78). Notably, Ran distribution can be more sensitive to cellular oxidative stress than is NLS-mediated classical nuclear import, particularly of small protein cargoes (73, 79). Moreover, oxidative stress increases the association between Ran and another nuclear export protein, chromosome region maintenance-1 (Crm1), and inhibits classical nuclear export (78). Therefore, in our study it is plausible that while mitochondrial dysfunction disrupts the Ran gradient and impairs nuclear export, the nuclear import of RFP-NLS remains relatively efficient. Overall, the precise defects in the NCT apparatus that are induced in disease pathogenesis depend on the neuronal subsets and molecular mechanisms involved in the neurodegenerative process. Future studies unraveling the effects of mitochondrial impairment on additional components of the NCT machinery, such as nuclear transport receptors, will lead to a better understanding of NCT disruption in DAergic neurons.

Our finding that pharmacological suppression of oxidative stress by a mitochondria-targeted antioxidant restored Nup153 and 107 levels in DAergic neural cells aligns with a prior observation of antioxidant treatments normalizing Nup153 in neural stem cells of an Alzheimer’s mouse model (61). Our observation reinforces the therapeutic and neuroprotective potential of mitochondrial antioxidants like mito-apocynin by intervening in key pathological mechanisms in PD.

Collectively, our data identify NPC and NCT dysregulation as potential mechanisms for mitochondrial dysfunction-induced DAergic neuron loss, which offers new mechanistic insights into PD pathogenesis. The pathologic alterations in Nups that we have uncovered using in vitro and in vivo models of mitochondrial impairment, as well as PD patient brains, may have extensive implications for neuronal function. Given that mitochondrial dysfunction is a common feature of several neurodegenerative diseases, targeting NPC-related impairment could be a promising strategy for therapeutic intervention of these disorders.

## Methods

Detailed description of the antibodies and chemicals used in this study is provided in supplemental methods.

### Cell culture and treatments

The rat-derived mesencephalic DAergic neural cell line, N27, was a kind gift from Dr. Kedar N. Prasad (University of Colorado Health Sciences Center, Denver, CO). N27 cells were cultured in RPMI 1640 medium (Sigma) supplemented with L-glutamine, 10% FBS (Sigma), and 100 units/mL penicillin and 100 µg/mL streptomycin (Thermo Fisher) and maintained at 37°C in a humidified atmosphere with 5% CO_2_ (80). N27 cells were treated with the specified concentrations (1 µM or 100 nM) of rotenone (Sigma) in serum-free RPMI media when the cells were 70-80% confluent. Concentrated rotenone stocks were prepared in DMSO (Sigma) and frozen at −80°C as single-use aliquots for up to 6 mo. An equal volume of DMSO was added to serum-free RPMI media in the control group and DMSO concentration was < 0.1% in all experiments to prevent toxicity. For hydrogen peroxide treatment, N27 cells were exposed to 30 µM hydrogen peroxide (Sigma) in complete RPMI media for 24 h (81). Mito-apocynin was synthesized as described previously (35) and dissolved in DMSO at a concentration of 50 mM. For rotenone and Mito-apocynin co-exposure, N27 cells were pre-incubated with 30 µM Mito-apocynin in serum-free media for 1 h before the addition of 1 µM rotenone and co-treatment for 6 h. TFAM-KD N27 cells were generated as described previously using lentivirus-based CRISPR/Cas9 TFAM knockout plasmids (35). We confirmed a 70% knockdown of TFAM in these cells by Western blot (Supp. Fig. 1A).

### Subcellular fractionation and Western blots

Nuclear and cytoplasmic fractions were extracted from cell pellets using NE-PER Nuclear and Cytoplasmic Extraction Reagents supplemented with Halt protease and phosphatase inhibitor cocktail (Thermo Fisher) following the manufacturer’s protocol. Protein was quantified using the Dye Reagent Concentrate (Bio-Rad) in a Bradford protein assay. We loaded 30 µg cytoplasmic and 20 µg nuclear fractions on Any kD Mini-PROTEAN TGX Precast Protein Gels (Bio-Rad) and ran at 100V for 1.5 h. The proteins were then transferred to nitrocellulose membranes (Bio-Rad) overnight at 25V. For normalization, total protein was stained using Revert 700 Total Protein Stain Kit (LI-COR), and the membranes were then blocked in Intercept (PBS) Blocking Buffer (LI-COR) for 1 h and incubated overnight in primary antibodies in blocking buffer with 0.1% Tween-20 (Bio-Rad) at 4°C. The following day, membranes were incubated in IR-dye-conjugated secondary antibodies (LI-COR) for 1 h and then scanned on either an Odyssey IR or Odyssey M imaging system (LI-COR). Western blot analysis was performed on Empiria Studio or Image Studio Lite software (LI-COR).

### Immunocytochemistry (ICC)

Immunofluorescent labeling of Nups in N27 cells was performed using a modified version of a previously published protocol (82). Briefly, N27 cells were cultured on poly D-Lysine-coated (Sigma) coverslips, treated with rotenone, and fixed in 2% paraformaldehyde (PFA) in PBS (EMS) for 10-15 min. The fixed cells were blocked and permeabilized for 1 h using 10 mg/mL BSA, 0.1% Triton X-100 (Sigma), and 0.02% SDS (Fisher) in PBS, and then incubated in primary antibodies diluted in blocking buffer for 1 h using gentle shaking. Samples were then incubated with Alexa-Fluor conjugated donkey secondary antibodies (Thermo Fisher) diluted in blocking buffer for 1 h, and nuclei were stained with 1:10,000 Hoechst dye (Thermo Fisher) in PBS for 8 min. F-Actin was stained with 1:1000 Phalloidin dye (Thermo Fisher) in PBS for 10 min. Coverslips were mounted on slides using Fluoromount (Sigma) mounting media. For immunofluorescent labeling of Ran, cells were fixed in 4% PFA and then blocked and permeabilized in 2% BSA, 0.5% Triton X-100, and 0.4% Tween-20 in PBS. Samples were incubated overnight at 4°C in primary antibody and all antibody dilutions were in 2% BSA.

### Generation of hiPSC-derived midbrain DAergic neurons, rotenone treatment and immunostaining

We obtained hiPSCs expressing mEGFP-tagged Nup153 (mEGFP-NUP153; AICS-0069 cl.88) that were developed at the Allen Institute for Cell Science (allencell.org/cell-catalog) and are available through Coriell (83). These hiPSCs are high-quality, certified fluorescently tagged cells in which mEGFP was monoallelically introduced at the N-terminus of Nup153 using CRISPR-Cas9 methodology. The hiPSCs were differentiated into floor-plate lineage DAergic neurons exactly as described in a previously published protocol (84), with two modifications. The cells were passaged twice using Accutase (Thermo Fisher), once on day 13 of differentiation at a ratio of 1:1.5 to avoid extreme overcrowding and detachment of neural progenitor cells, and again between days 20 and 25 of differentiation onto fresh Matrigel-coated (Corning) plates. Secondly, 100 units/mL penicillin and 100 µg/mL streptomycin were added to all basal media. Supp. Fig. 4 shows the expression of the pluripotency marker in hiPSCs, neural progenitor markers at day 14 of differentiation, and neuronal markers at day 35 of differentiation. Between days 35-40 of differentiation, the DAergic neurons were treated with 100 µM rotenone in complete differentiation media for 24 h. Concentrated rotenone stocks were used and an equal volume of DMSO was added to the control group, as described above for N27 cell treatment. For all immunostaining experiments, cells were fixed using 4% PFA on Matrigel-coated coverslips, and then blocked, permeabilized and incubated with antibodies as described above for Ran ICC in N27 cells.

### Immunohistochemistry (IHC) of paraffin brain sections from MitoPark mice

MitoPark mice were kindly provided by Dr. Nils-Goran Larson at the Karolinska Institute, Stockholm (36). The MitoPark and littermate control (C57BL/6) mice used in this study were bred, genotyped, and extensively characterized at UGA and ISU. These mice were fed and maintained under standard conditions as previously described (40). Equal numbers of age- and sex-matched MitoPark and littermate control mice were transcardially perfused with PFA as described previously (85), and formalin fixed brains were sent to the Histology lab at the UGA College of Veterinary Medicine for paraffin embedding and sectioning. The brain sections were 5 microns in thickness. The immunofluorescent labeling protocol used in this study was adapted from a previously published protocol (48). Briefly, the slides were deparaffinated by heating to 59°C for up to 5 min on a slide warmer, followed by sequential incubation in xylene x 2 (Thermo Fisher), 100%, 90% and 70% ethanol (Thermo Fisher) for 5 min each, and rehydrated in distilled water. Antigen retrieval was performed by steaming the slides in IHC-Tek epitope retrieval solution (IHC World) for 1 h. The tissue sections were permeabilized in 0.4% Triton X-100 in PBS for 8 mins and blocked with 10% normal donkey serum (Sigma) and 0.2% Triton X-100 in PBS for 1 h. The slides were then incubated with primary antibodies in 10% normal donkey serum overnight at 4°C in humidity chamber. The following day, sections were incubated with Alexa-Fluor conjugated secondary antibodies in 3% normal donkey serum for 1 h, nuclei were stained with a 1:10,000 diulation of Hoechst dye for 8 mins, and slides were mounted with Fluoromount.

### Human post-mortem brain tissues

We obtained freshly frozen SN tissue samples from the brain banks at Miller School of Medicine, University of Miami, FL, and fresh frozen SN cryostat sections from the Banner Sun Health Research Institute, AZ. For Western blot, SN tissue samples from PD patients and healthy controls were lysed and homogenized in RIPA buffer (Thermo Fisher) supplemented with Halt protease and phosphatase inhibitor cocktail and 1:100 PMSF (Sigma). The lysates were sonicated and centrifuged, and then 15-30 µg protein was used to perform Western blot as described above. Cryostat sections were used for immunostaining of Nups. The samples were thawed to room temperature, washed thrice for 10 mins in PBS, and blocked for 1 h with 5% donkey serum, 0.05% Triton X-100. Slides were incubated overnight at 4°C with primary antibodies diluted in 1:5 diluted blocking buffer (1% donkey serum, 0.01% Triton X-100). The following day, sections were washed 6 x 5 mins in PBS and incubated with Alexa-Fluor conjugated secondary antibodies in diluted blocking buffer. The slides were again washed 6 x 5 mins in PBS, incubated with 1:10,000 Hoechst dye for 5 min, washed and allowed to dry overnight. The next day, the sections were washed in PBS for 3 mins and then sequentially dehydrated in 50%, 70%, 95% and 100% ethanol for 1 min each, dipped twice in xylene, and coverslips were mounted on the slides using Fluoromount.

### Nucleocytoplasmic transport assay

N27 cells were transduced with pLVX-EF1alpha-2xGFP:NES-IRES-2xRFP:NLS, which was a gift from Fred Gage (Addgene plasmid # 71396 ; http://n2t.net/addgene:71396 ; RRID:Addgene_71396) (49). This is a lentiviral mammalian expression plasmid consisting of GFP and RFP attached to nuclear export and localization signals (NES:GFP and NLS:RFP), respectively, with an internal ribosome entry site between them. The cells were incubated with transduction media for 72 h and then exposed to 1 µM rotenone in fresh serum-free media for 6 h. The cells were then fixed in 4% PFA for 10 mins and mounted in Fluoromount. Fluorescence microscopy and analysis were performed on these slides as described below.

### Super-resolution structured illumination, confocal, and fluorescence microscopy

Super-resolution structured illumination (SR-SIM) imaging was performed with 3 grid rotations and optimal z sectioning using the 100x objective on the Zeiss ELYRA S1 microscope. SIM processing was performed using automatic parameters on ZEN 2011 (black edition) software. For each ICC experiment of Nups in N27 cells, at least 50 images of control and treated cells were captured and processed using identical parameters, with each image containing a single cell. For Nups 153, 107 and 62, single z-slice images showing Nup staining at the nuclear boundary are presented. The fluorescence signal for these Nups was also quantified from single z-slice images using a published Cell Profiler ver. 4.2.4 pipeline (86) with some optimizations of thresholding and object size range to accurately identify nuclei and Nup signal. To identify Nup signal objects, Otsu thresholding was used and integrated intensity was measured for Nup objects at the nuclear envelope only. The fluorescence signal for Nups 50 and 214 was not resolved at the nuclear boundary, so maximum intensity projections of z slices are presented for them. The integrated intensity of fluorescence signal for these Nups was also quantified from maximum intensity projection images using Cell Profiler. In this pipeline, Nup signal objects were identified from within a nuclear mask using Otsu thresholding with a threshold smoothing scale of 1.3488, a correction factor of 1, lower and upper bounds of 0.05 and 1, and no declumping. Cell Profiler subplots of Nup objects that were quantified are shown in Fig. 2. SR-SIM imaging and processing of Nup153-EGFP in hiPSC-derived DAergic neurons were also performed using the conditions described above.

Confocal imaging of immunofluorescently labeled Nups in MP SN sections, human SN sections and Ran in N27 cells was performed using the 100x objective and standard imaging conditions on the Zeiss LSM880 upright confocal microscope and ZEN black software. Both SR-SIM and confocal microscopy were performed at the Biomedical Microscopy Core at UGA.

Fluorescence microscopy of immunostained MP and human SN sections, Ran in cells and tissue sections, hiPSC-derived DAergic neurons, and reporter constructs in the nucleocytoplasmic transport assay was performed using the Keyence BZ-X810 microscope. The processing and quantification of these fluorescence images were performed on the Keyence Analyzer software. To quantify the fluorescence signal from the nuclei, the single extraction module was used to extract target protein signal from the nuclear mask constructed over Hoechst signal. Average brightness integration values for the specific channel were measured. To calculate the nuclear-to-cytoplasmic ratios of Ran GTPase and GFP and RFP reporters, the nuclear signal was subtracted from the total signal for each channel to get the cytoplasmic signal value, which was then used to calculate the ratio. To quantify Nup62 cytoplasmic puncta in N27 cells treated with 100 nM rotenone, the threshold was manually set to identify each Nup62 speck separately, nuclear objects were deselected, and the number of cytoplasmic objects was divided by the total number of cells per image to get a count of the Nup62 puncta/cell.

### Statistics

Statistical analyses of the data were performed using GraphPad Prism version 9 and 10 (GraphPad). Statistics were performed based on the assumption of normality using a two-tailed unpaired student’s t-test with Welch’s correction for unequal variances (2 groups), an ordinary one-way ANOVA with Tukey’s multiple comparisons test (3 groups), and multiple unpaired t-tests with Holm-Sidak method to correct for multiple comparisons (grouped data; data from each treatment group was compared to its control). All data is presented as mean ± SEM. The threshold for statistical significance is p < 0.05. For all experiments, the number of independent repeats or biological samples (mouse and human samples) was at least N= 3. Datasets with several data points, such as those from microscopy analyses, are shown as dot plots to show the full data distribution and variation, and bar graphs with individual data points are used to display smaller datasets. The specific number of independent experiments or sample number, the statistical test used for data comparison, and the statistical significance of data are stated in the legend for each figure panel.

### Data availability

All data are available in the main text, supplemental materials, or from the authors upon request. Values for all data points in graphs are reported in the Supporting Data Values file.

## Credit Author Contributions Statement

**Zainab Riaz**: Conceptualization, Investigation, Methodology, Data curation, Visualization, Validation, Formal Analysis, Writing—original draft, Writing—review and editing. **Yuan-Teng Chang**: Data curation, Investigation, Methodology, Validation. **Gary Zenitsky**: Writing—review and editing. **Huajun Jin**: Formal analysis, Writing—review and editing. **Vellareddy Anantharam**: Conceptualization, Supervision, Project Administration Resources, Software. **Arthi Kanthasamy**: Conceptualization, Funding Acquisition, Resources. **Anumantha Kanthasamy**: Conceptualization, Funding Acquisition, Project Administration, Resources, Supervision, Writing—review and editing.

