## Supplementary document for "Mitochondrial dysfunction Impairs the Nuclear Pore Complex in Parkinson’s Disease Pathogenesis"

### Supplemental materials

**Supplemental methods:**

***Antibodies***

Supp. Table 1. List of antibodies used in the study.

| Antibodies | Source | Catalog no. | Application |
|---|---|---|---|
| Rabbit Anti-Nup50 | Abcam | ab137092 | Western blot (1:2000)<br>ICC (1:500 – 1:2000) |
| Rabbit Anti-Nup50 | Abcam | ab85915 | IHC (1:250) |
| Mouse Anti-Nup62 | BD Biosciences | 610497 | Western blot (1:2000)<br>ICC (1:500 – 1:2000) |
| Rabbit Anti-Nup62 | Thermo Fisher Scientific | PA5-21882 | IHC (1:100) |
| Rabbit Anti-Nup107 | Abcam | ab73290 | Western blot (1:2000) |
| Mouse Anti-Nup107 | Thermo Fisher Scientific | MA1-10031 | ICC (1:500)<br>IHC (1:50) |
| Mouse Anti-Nup153 | Abcam | ab96462 | Western blot (1:1000)<br>ICC (1:500- 1:2000)<br>IHC (1:100) |
| Rabbit Anti-Nup214 | Abcam | ab70497 | ICC (1:500) |
| Mouse Anti- Nup358 | Thermo Fisher Scientific | MA1-847 | ICC (1:500) |
| Mouse Anti-Ran | BD Biosciences | 610341 | ICC (1:500)<br>IHC (1:100) |
| Rabbit Anti-GAPDH | Cell Signaling | 2118 | Western blot 1:10,000 |
| Mouse Anti-TFIID | Santa Cruz | sc-56794 | Western blot 1:500 |
| Mouse Anti-414 | Abcam | ab24609 | IHC (1:100 – 1:500) |
| Rabbit Anti-Tyrosine Hydroxylase | Millipore Sigma | AB152 | ICC and IHC (1:2000 – 1:5000) |
| Mouse Anti-Tyrosine Hydroxylase | Millipore Sigma | MAB318 | IHC (1:2000 – 1:5000) |
| Mouse Anti-Ubiquitin | Thermo Fisher Scientific | 13-1600 | IHC (1:100) |
| Rabbit Anti-G3BP1 | Thermo Fisher Scientific | PA5-29455 | ICC (1:500) |
| Mouse Anti-LaminB1 | Santa Cruz Biotechnology | sc374015 | ICC (1:500) |
| Mouse Anti-Oct4 | Stem Cell Technologies | 60093 | ICC (1:500) |
| Mouse Anti-Nestin | Stem Cell Technologies | 60091 | ICC (1:500) |
| Rabbit Anit-Pax6 | BioLegend | 901301 | ICC (1:500) |

***Chemicals***

Supp. Table 2. List of chemicals and reagents used in the study.

| Chemical | Source | Catalog no. |
|---|---|---|
| Accutase | Thermo Fisher Scientific | 00-4555-56 |
| Alexa Fluor 488 Phalloidin | Thermo Fisher Scientific | A12379 |

| | | |
|---|---|---|
| Ascorbic acid | Sigma | A4034 |
| B27 supplement | Thermo Fisher Scientific | 17504044 |
| BDNF | R&D Systems | 248-BD-025 |
| Beta mercaptoethanol | Sigma | M3148 |
| Bio-Rad Protein Assay Dye Reagent Concentrate | Bio-Rad | 5000006 |
| Bovine serum albumin (BSA) | Sigma | A9647 |
| CHIR99021 | Stemgent | 04-0004 |
| D- Glucose solution | Thermo Fisher Scientific | A2494001 |
| DAPT | Tocris | 2634 |
| dcAMP | Sigma | D0627 |
| Dimethyl sulfoxide (DMSO) | Sigma | D8418 |
| DMEM/F12, Glutamax | Thermo Fisher Scientific | 10565-018 |
| Donkey serum | Sigma | D9663 |
| Ethanol | Thermo Fisher Scientific | T032021000 |
| Fetal Bovine Serum (FBS) | Sigma | 12306C |
| FGF-8a | R&D Systems | 4745-F8-050 |
| Fluoromount mounting medium | Sigma | F4680 |
| GDNF | R&D Systems | 212-GD-050 |
| Glutamax | Thermo Fisher Scientific | 35050-061 |
| Hoechst 33342 | Thermo Fisher Scientific | H1399 |
| Hydrogen peroxide | Sigma | H1009 |
| IHC-Tek Epitope Retrieval Solution | IHC World | IW-1100 |
| Intercept (PBS) Blocking Buffer | LI-COR | 927-70001 |
| Knockout DMEM/F12 | Thermo Fisher Scientific | 12660 |
| Knockout Serum | Thermo Fisher Scientific | 10828 |
| LDN193189 | Stemgent | 04-0074 |
| Matrigel | Corning | 356231 |
| MitoTracker Red CMXRos | Thermo Fisher Scientific | M7512 |
| N2 supplement | Thermo Fisher Scientific | 17502-048 |
| NE-PER Nuclear and Cytoplasmic Extraction Reagents | Thermo Fisher Scientific | 78833 |
| Neurobasal media | Thermo Fisher Scientific | 21103-049 |
| Non-essential amino acids | Thermo Fisher Scientific | M7145 |
| Paraformaldehyde (PFA) | Electron Microscopy Sciences | 19210 |
| Penicillin-Streptomycin | Thermo Fisher Scientific | 15140122 |
| Phenylmethanesulfonyl fluoride (PMSF) | Sigma | 93482 |
| Phosphate buffered saline (PBS) | Fisher Scientific | BP661-10 |
| Poly D-Lysine | Sigma | P6407 |
| Puromorphamine | Stemgent | 04-0009 |

| | | |
|---|---|---|
| Revert 700 Total Protein Stain Kit | LI-COR | 926-11010 |
| RIPA buffer | Thermo Fisher Scientific | 89901 |
| Rotenone | Sigma | R8875 |
| RPMI 1640 | Sigma | R8758 |
| SB431542 | Cayman Chemical | 13031 |
| Shh (C24II) | R&D Systems | 1845-SH-100/CF |
| Sodium dodecyl sulfate (SDS) | Fisher Scientific | BP166-100 |
| TGFβ3 | R&D Systems | 243-B3-002 |
| Triton X-100 | Sigma | T8787 |
| Tween-20 | Bio-Rad | 1610781 |
| Xylene | Thermo Fisher Scientific | A11358.0F |
| SYTOX™ Green Nucleic Acid Stain | Thermo Fisher Scientific | S7020 |

***Primers***

Supp. Table 3. List of primers used in the study.

| Primers | Source | Assay ID |
|---|---|---|
| Nup107 | IDT | Rn.PT.58.11400635 |
| Nup153 | IDT | Rn.PT.58.46109978 |
| Nup50 | IDT | Rn.PT.58.11562548 |
| Nup62 | IDT | Rn.PT.58.34451535 |
| Nup214 | IDT | Rn.PT.58.37919624 |

***Real-time RT-qPCR***

Total RNA was isolated from N27 cells with Trizol. RNA was reverse transcribed to synthesis cDNA using High-Capacity cDNA Reverse Transcription Kit according to the manufacturer's protocol. A total of 1 µl of cDNA was added into a 10 µl reaction and amplified with PowerUp SYBR Green Master Mix for qPCR. All primers for qPCR were purchased from IDT (Supplementary Table 3).

***SYTOX Green Assay***

N27 cells were incubated with 1 µM of SYTOX Green staining solution in the last 30 min of the rotenone treatment time. The cells were washed 3 times with a phosphate-free buffer and then fixed with 4% PFA for 15 min. After fixation, cells were washed 2 more times, counterstained with Hoechst, and mounted in Fluoromount.

***Transmission electron microscopy***

N27 cells were treated with 1 µM rotenone in serum-free RPMI media for 6 h, harvested, and pelleted by spinning at 300 x g for 5 min. After thoroughly washing the cell pellet with PBS, the cells were centrifuged again and then fixed overnight with a solution containing 2% glutaraldehyde and 2% paraformaldehyde in 0.1M Cacodylate-HCl buffer, pH 7.25 at 4°C. After being washed several times in 0.1M Cacodylate-HCl buffer, cells were agar-enrobed with 3% Noble Agar at a temperature of 60°C. Once cooled, agar-cell pellets were extracted from Eppendorf tubes and placed in 0.1M Cacodylate-HCl buffer to continue processing. Cells were post-fixed with 1%

buffered osmium for 1 h and rinsed in four changes of deionized water. The cells were dehydrated in increasing concentrations of ethanol (30%, 50%, 75%, 95%, 100%), then infiltrated with propylene oxide, and then infiltration of propylene oxide and increasing amounts of Epon-Araldite plastic (1). After several long changes in 100% Epon-Araldite plastic, the cell pellets were embedded in fresh Epon-Araldite plastic and allowed to polymerize in a 60°C oven for 3 days. Ultrathin sections were cut on a Reichert Ultracut S ultramicrotome, placed on clean 200-mesh Cu Hex grids, and stained with 2% aqueous uranyl acetate and Reynold's lead citrate (2). Sections were examined with a JEOL JEM-1011 Transmission Electron Microscope (JEOL USA, Inc.) at an accelerating voltage of 100 kV. Digital images were acquired using an AMT Imaging System (Advanced Microscopy Techniques). All processing, sectioning, and imaging was performed at the Georgia Electron Microscopy core facility at the University of Georgia campus.

##### *MitoTracker staining*

N27 cells were incubated with 50nM MitoTracker Red dye in pre-warmed PBS for 10 min following treatments with rotenone and Mito-apocynin. Cells were washed in PBS after staining, counterstained with Hoechst, and mounted in Fluoromount.

##### *References to supplemental methods*

1. Mollenhauer HH. Plastic Embedding Mixtures for Use in Electron Microscopy *Stain Technol.* 1964;39:111-4.
2. Reynolds ES. The use of lead citrate at high pH as an electron-opaque stain in electron microscopy. *J Cell Biol.* 1963;17(1):208-12.

**Supplemental figures:**

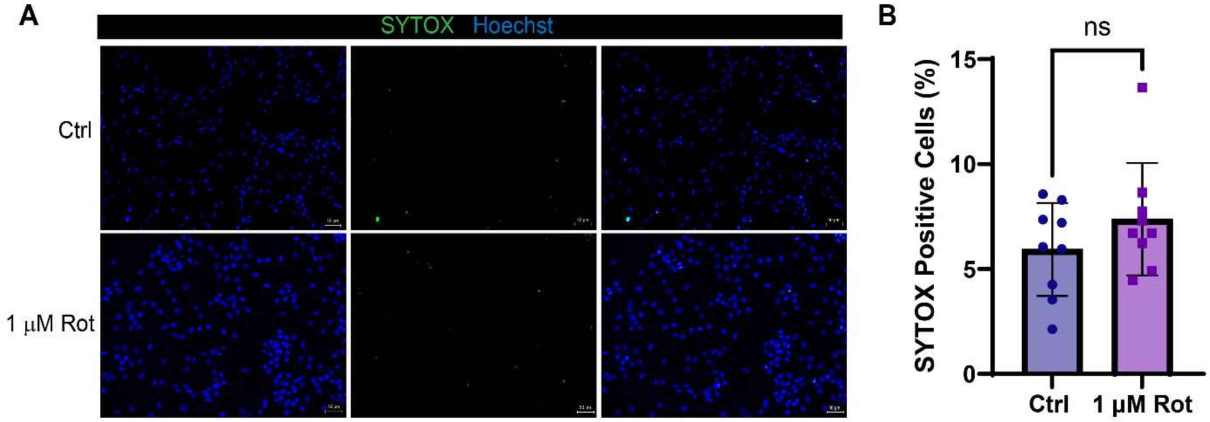


**Supp. Fig. 1 Rotenone treatment does not induce significant cell death in N27 DAergic neural cells.**

(A) Representative images of SYTOX green assay of N27 cells treated with 1 μM rotenone for 6 h. (B) Quantification of SYTOX positive cells as a percentage of the total number of cells that stained by Hoechst. Data from 3 independent experiments shown as mean ± SD. Unpaired t-test. ns, non-significant. Scale bar: 50 μm.

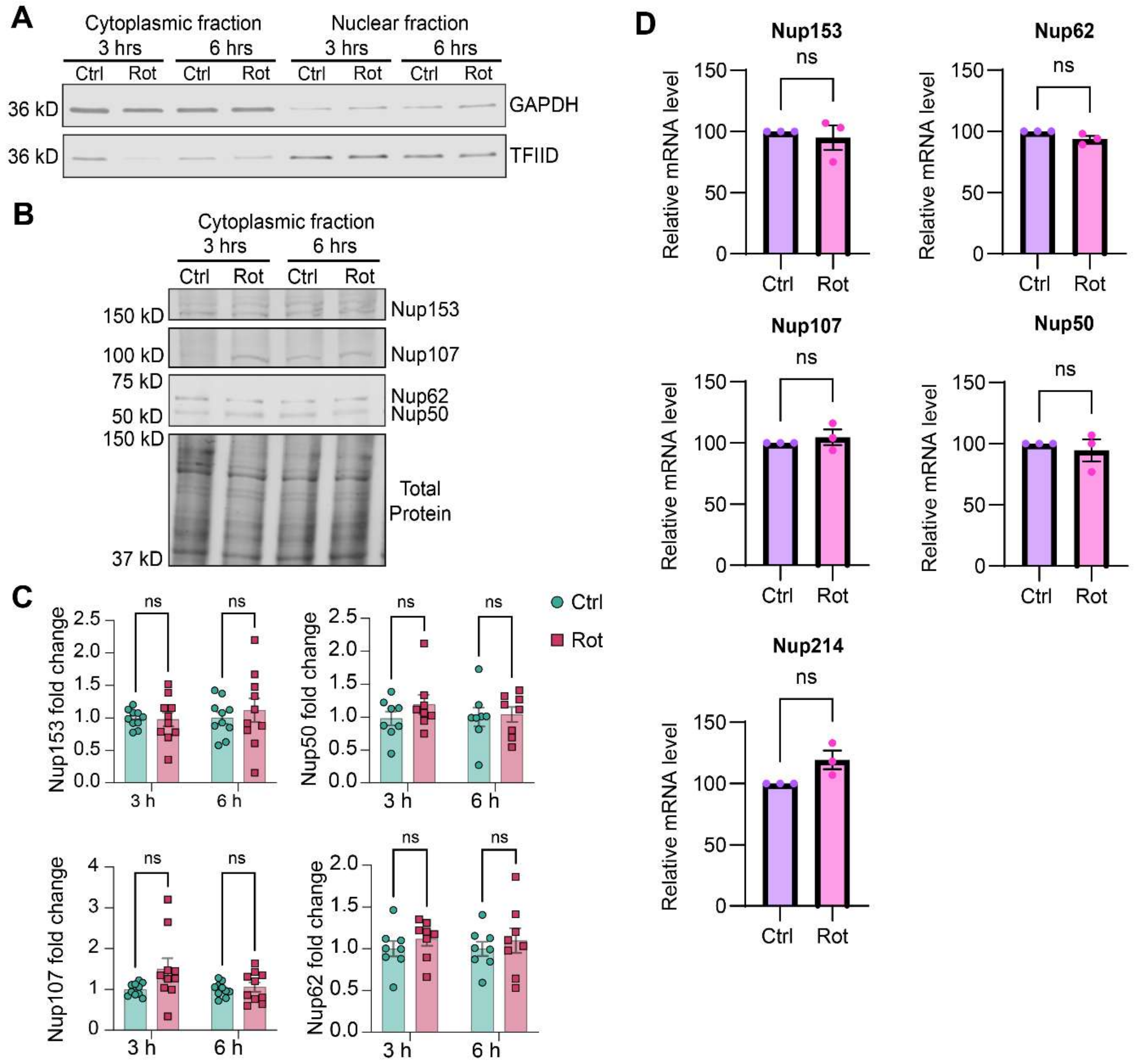


**Supp. Fig. 2 Nups are not mislocalized to the cytoplasm and their mRNA levels are unchanged in N27 DAergic neural cells under acute mitochondrial stress.**

(A) Representative Western blot image of cytoplasmic and nuclear fractions from control and rotenone exposed N27 cells for the cytoplasmic marker, GAPDH, and nuclear marker, TFIID. (B) Representative Western blot images of Nups 153, 107, 50 and 62 in cytoplasmic fraction of N27 DAergic neural cells exposed to 1μM rotenone for 3 and 6 h. (C) Quantification of Nups from the cytoplasmic fraction from at least 4 independent experiments, with 2 technical repeats each. Multiple unpaired t-tests with Holm-Sidak method to correct for multiple comparisons. ns, not significant. All western blots were normalized to total protein. (D) qRT-qPCR for Nups 153, 107, 62, 50, and 214 in N27 cells exposed to 1μM rotenone for 6 h. Data from 3 independent experiments and calculated using ΔΔCt method, normalized with GAPDH. Unpaired t-tests with Welch's correction for unequal variances. ns, not significant. All data are shown as fold change relative to control (mean ± SEM).

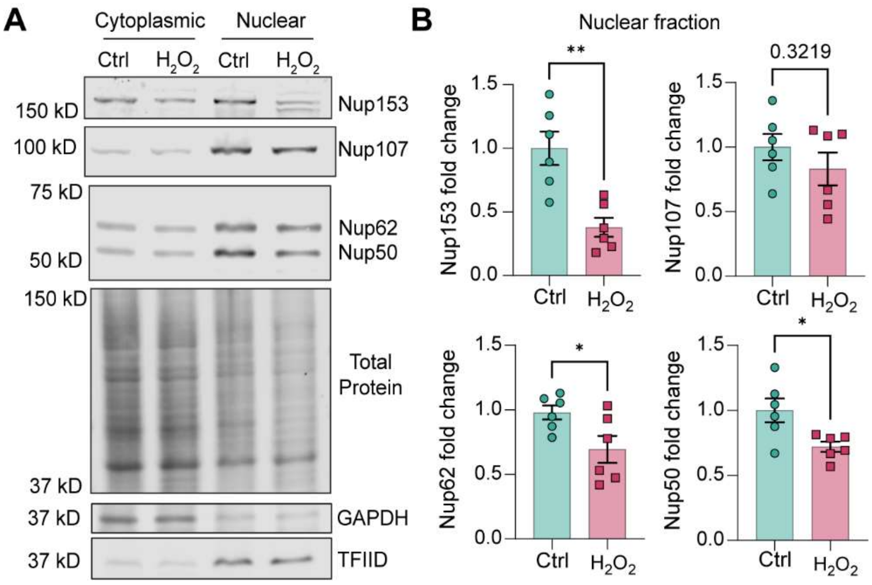


**Supp. Fig. 3 Hydrogen peroxide-induced oxidative stress downregulates Nups in the nuclear fraction of N27 DAergic neural cells.**

(A) Representative western blot images of Nups 153, 107, 62 and 50 in cytoplasmic and nuclear fractions of control and hydrogen peroxide ($H_2O_2$) exposed N27 cells. (B) Quantification of Nups in the nuclear fraction from 3 independent experiments, with 2 technical repeats each. Unpaired t-tests, $^*p<0.05$; $^{**}p<0.01$, p-value shown. All western blots were normalized to total protein and Nup levels are shown as fold change relative to control (mean ± SEM).

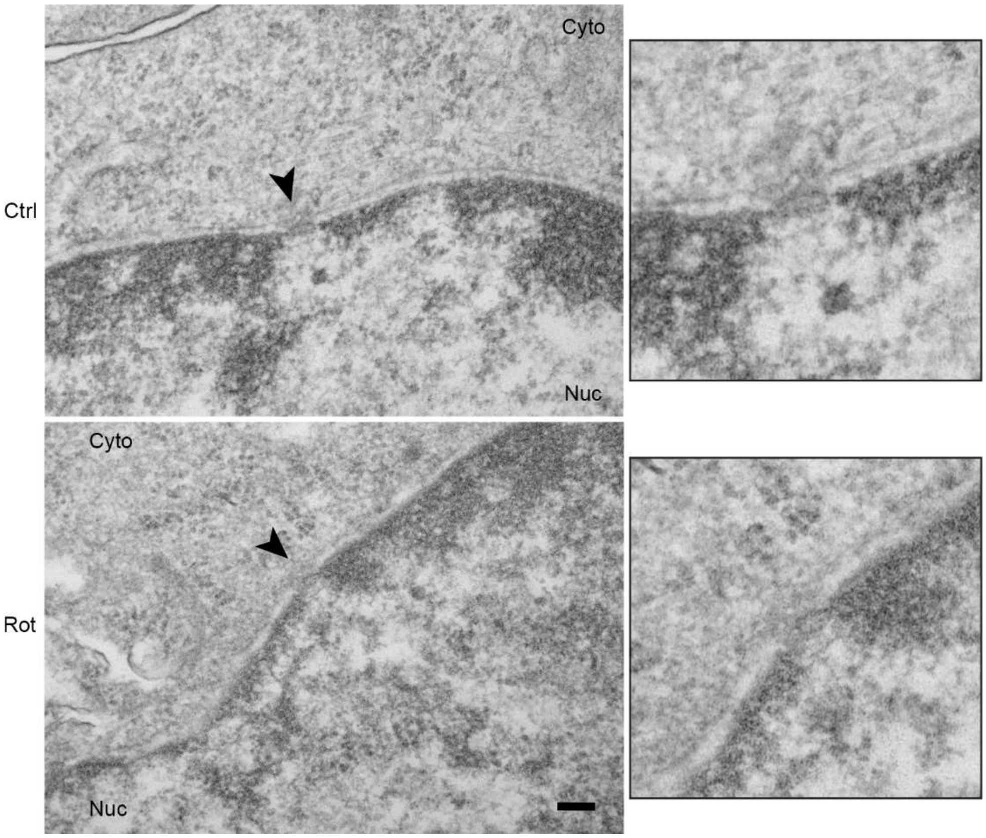


**Supp. Fig. 4 Transmission electron microscopy of NPC morphology in control and rotenone exposed N27 DAergic neural cells.**

Cross-sectioned nuclear membrane of N27 cells exposed to 1μM rotenone for 6 h, compared to control. The cytoplasmic and nuclear sides are marked as “cyto” and “nuc” respectively. Regions of NPCs are marked by arrows. Zoomed insets show twofold-enlarged images of NPCs. Imaged at 20000x. Scale bars, 100 nm.

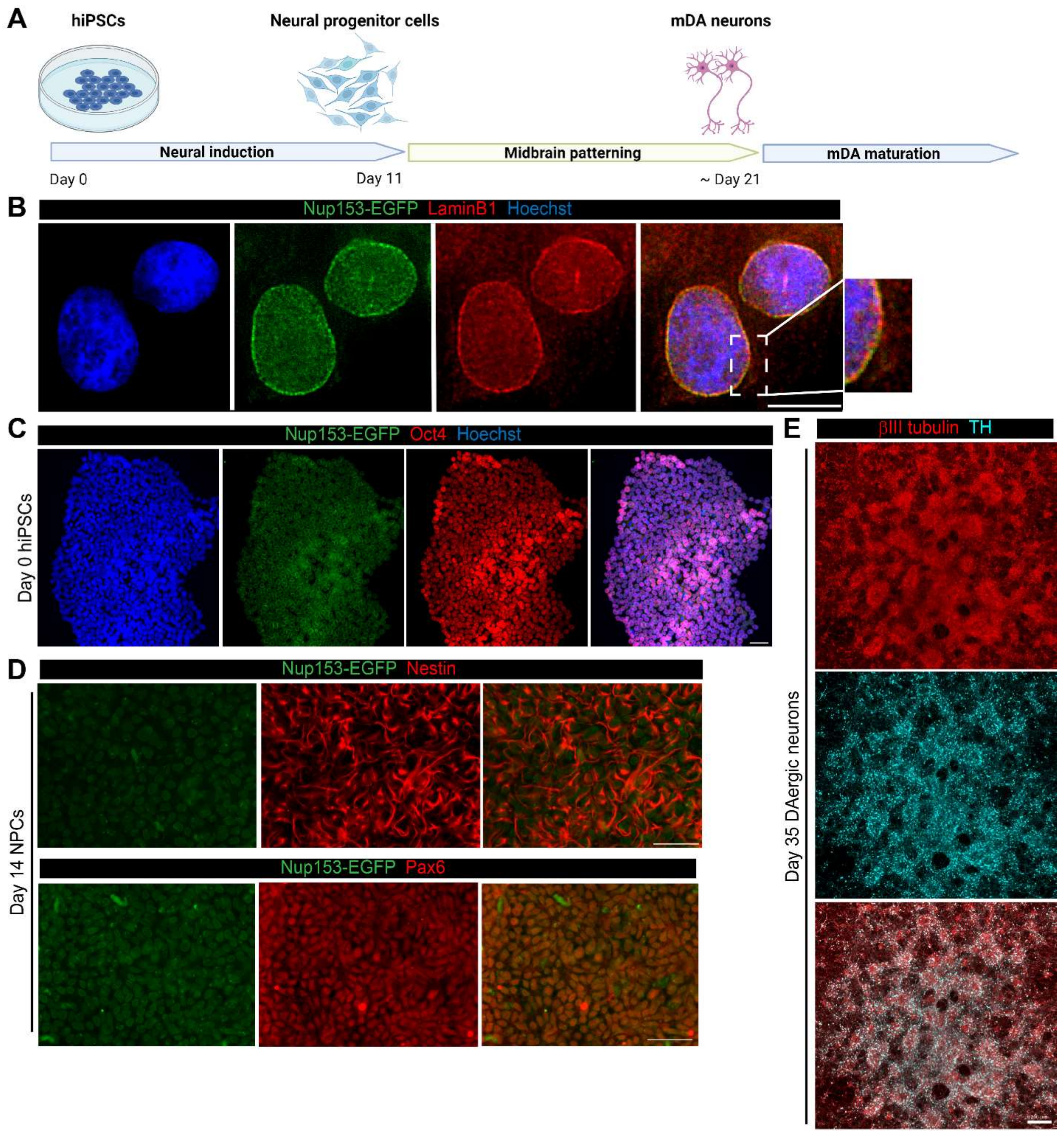


**Supp. Fig. 5 Differentiation of Nup153-EGFP tagged hiPSCs into midbrain DAergic neurons.**

(A) Schematic overview of the midbrain DAergic neuron differentiation workflow. (B) Immunocytochemistry analysis of hiPSCs showing colocalization of Nup153-EGFP with LaminB1 at the nuclear membrane. Scale bars, 10 µm. (C) Immunofluorescence images of day 0 hiPSCs showing Nup153-EGFP expression and stained for the pluripotency marker, Oct4. Scale bars, 50 µm. (D) Immunocytochemistry of day 14 Nestin and Pax6-positive neural progenitor cells. Scale bars, 50 µm. (E) Immunofluorecence images showing βIII tubulin and TH -positive midbrain neurons on day 35 of differentiation. Scale bar, 200 µm.

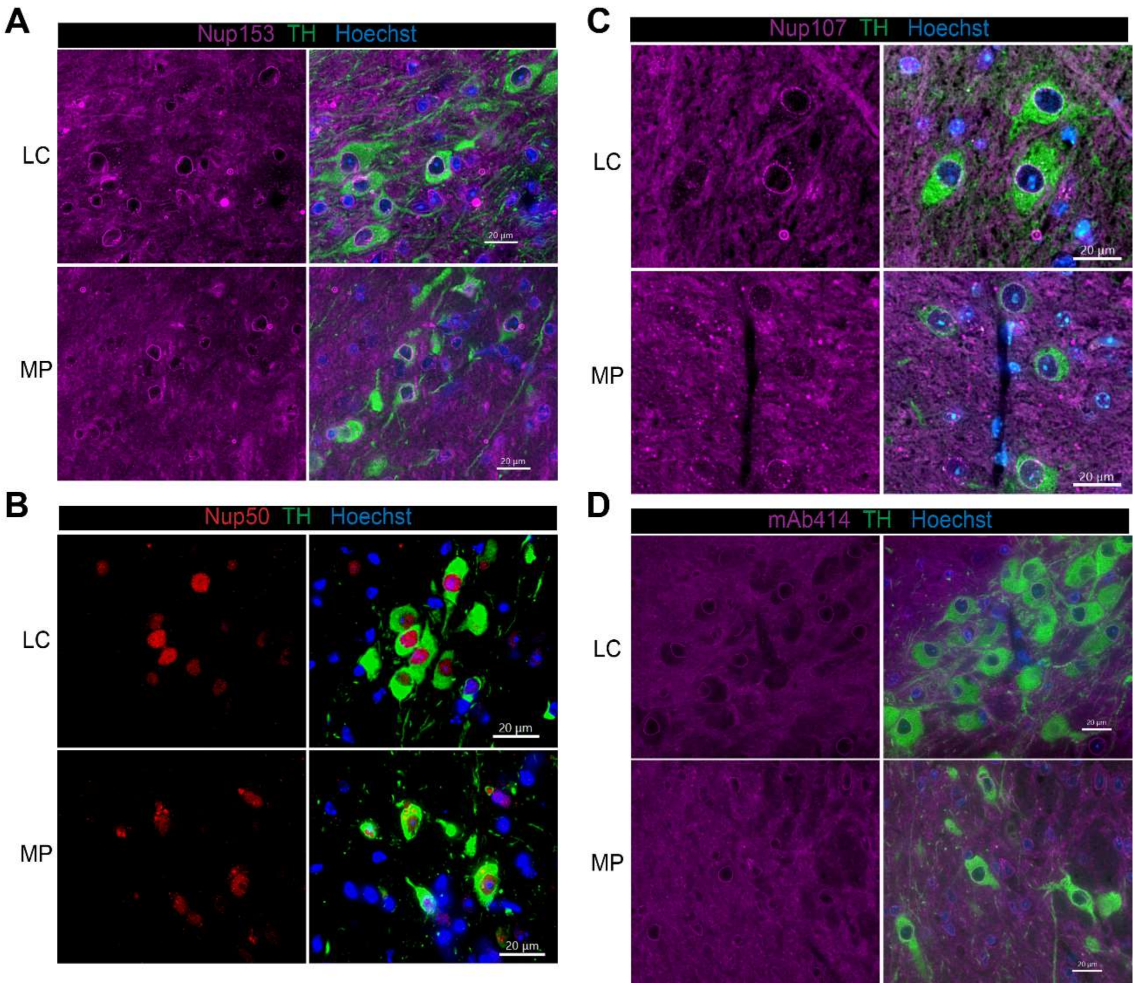


**Supp. Fig. 6 Fluorescence microscopy images of Nups in substantia nigra of littermate control and MitoPark transgenic mice.**

Representative fluorescence microscopy images of SN sections of MitpPark (MP) and littermate control (LC) mice immunostained for TH and (A) Nup153, (B) Nup50, (C) Nup107 and (D) NPC stained by the mAb414 antibody. Quantification of fluorescence signal integrated intensity from these images shown in Fig. 4. Scale bars, 20 μm.

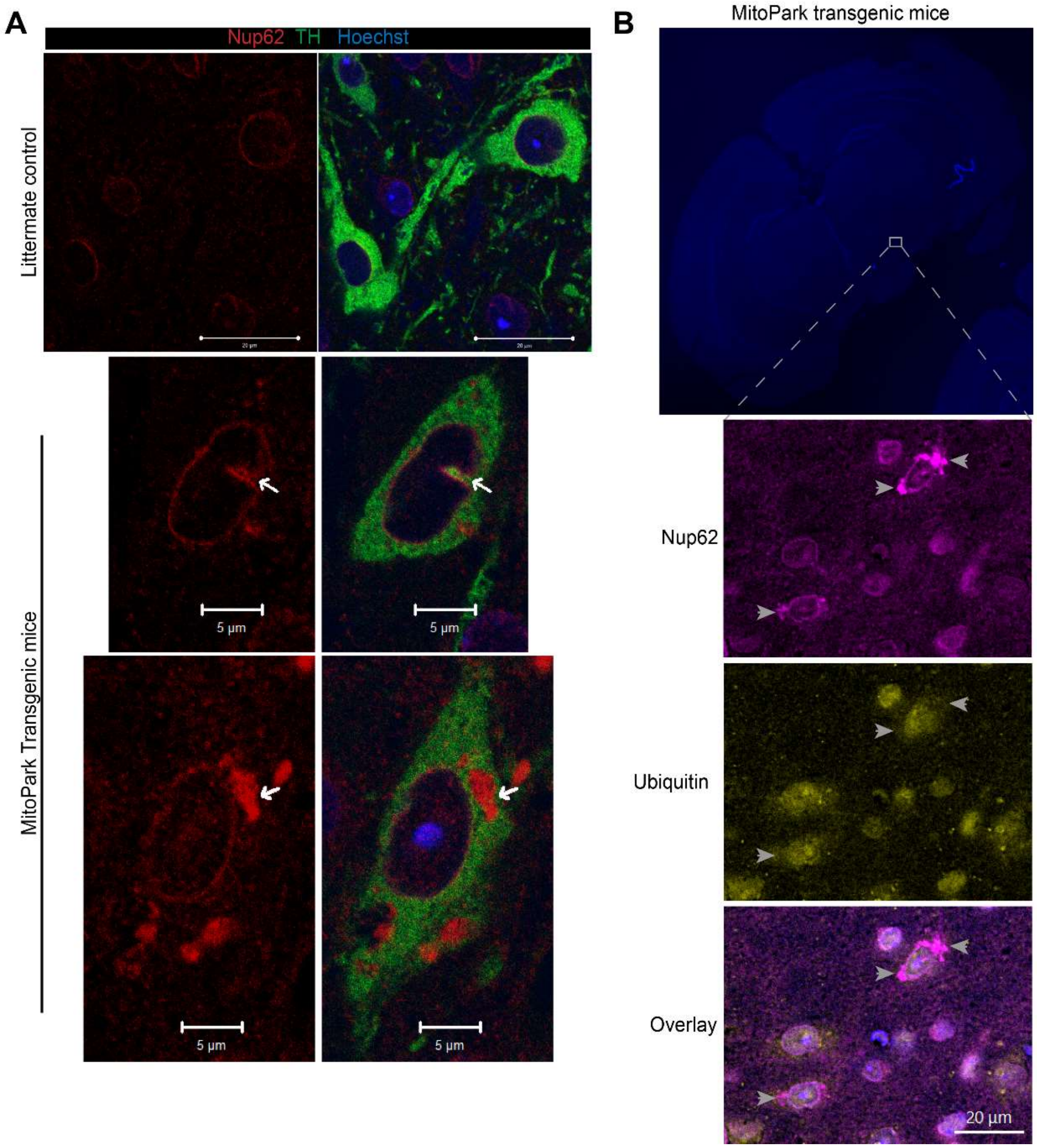


**Supp. Fig. 7 Nup62 is mislocalized to extranuclear aggregates that lack ubiquitin in nigral DAergic neurons of MitoPark mice.**

(A) Representative confocal images of SN sections of MitoPark and littermate control mice immunostained for TH and Nup62 from different fields. Arrows show nuclear membrane invagination and extranuclear aggregates of Nup62. Scale bars, 20 µm in littermate control panel; 5 µm in MitoPark mice panel. (B) Representative images of SN sections of MitoPark mice immunostained for Nup62 and ubiquitin. Arrows show extranuclear Nup62 and lack of ubiquitin and Nup62 signal colocalization. Scale bar, 20 µm.

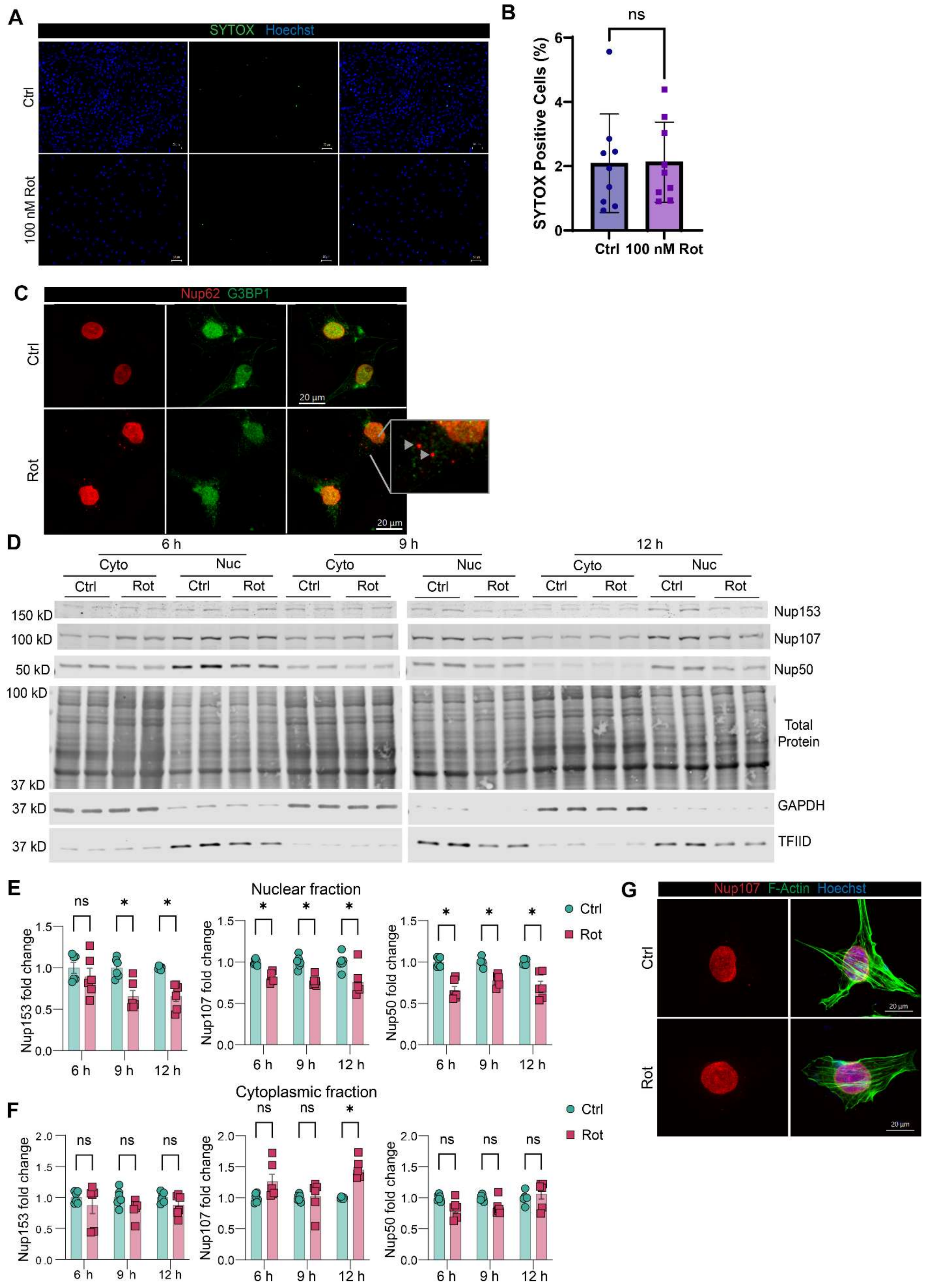

A
SYTOX Hoechst
Ctrl
100 nM Rot
B
ns
SYTOX Positive Cells (%)
Ctrl
100 nM Rot
C
Nup62 G3BP1
Ctrl
Rot
20 µm
D
6 h
9 h
12 h
Cyto
Nuc
Ctrl
Rot
150 kD
100 kD
50 kD
37 kD
Nup153
Nup107
Nup50
Total Protein
GAPDH
TFIID
E
Nuclear fraction
Nup153 fold change
Nup107 fold change
Nup50 fold change
6 h
9 h
12 h
Ctrl
Rot
F
Cytoplasmic fraction
G
Nup107 F-Actin Hoechst

**Supp. Fig. 8 Nups are depleted in the nuclei and Nup62 is mislocalized to the cytoplasm in N27 DAergic neural cells exposed to low-dose rotenone.**

(A) Representative images of SYTOX green viability assay of N27 cells treated with 100 nM rotenone for 12 h. (B) Quantification of SYTOX positive cells as a percentage of the total number of cells that stained by Hoechst. Scale bar: 50 μm. Data from 3 independent experiments shown as mean ± SD. Unpaired t-test. (C) Representative immunofluorescence images of Nup62 and the stress granule marker, G3BP1, in N27 cells exposed to 100nM rotenone for 12 h. Arrows show extranuclear Nup62 aggregates and lack of G3BP1 and Nup62 signal colocalization. Scale bars, 20 μm. (D) Representative western blot images of Nup153, 107, and 50 in cytoplasmic and nuclear fractions of N27 cells exposed to 100 nM rotenone for 6, 9 and 12 h. Quantification of Nups from the (E) nuclear and (F) cytoplasmic fractions from 3 independent experiments, with 2 technical repeats each. Multiple unpaired t-tests with Holm-Sidak method to correct for multiple comparisons. (G) Representative immunofluorescence images of Nup107 in N27 cells exposed to 100 nM rotenone for 12 h. Scale bars, 20 μm. *$p<0.05$; ns, not significant.

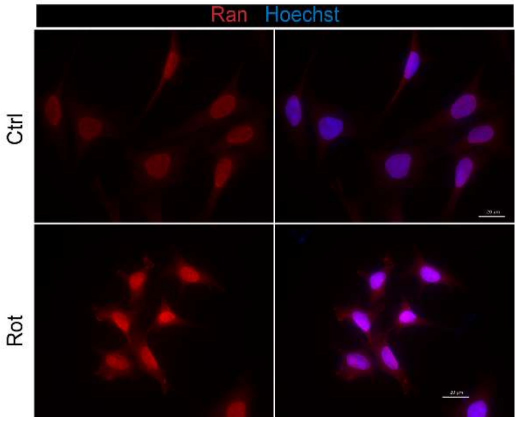


**Supp. Fig. 9 Mitochondrial dysfunction disrupts the Ran gradient in N27 DAergic neural cells.**

Representative fluorescence microscopy images of Ran distribution in control and rotenone exposed N27 cells. Quantification of nuclear/ cytoplasmic Ran distribution from these images is shown in Fig. 7C.

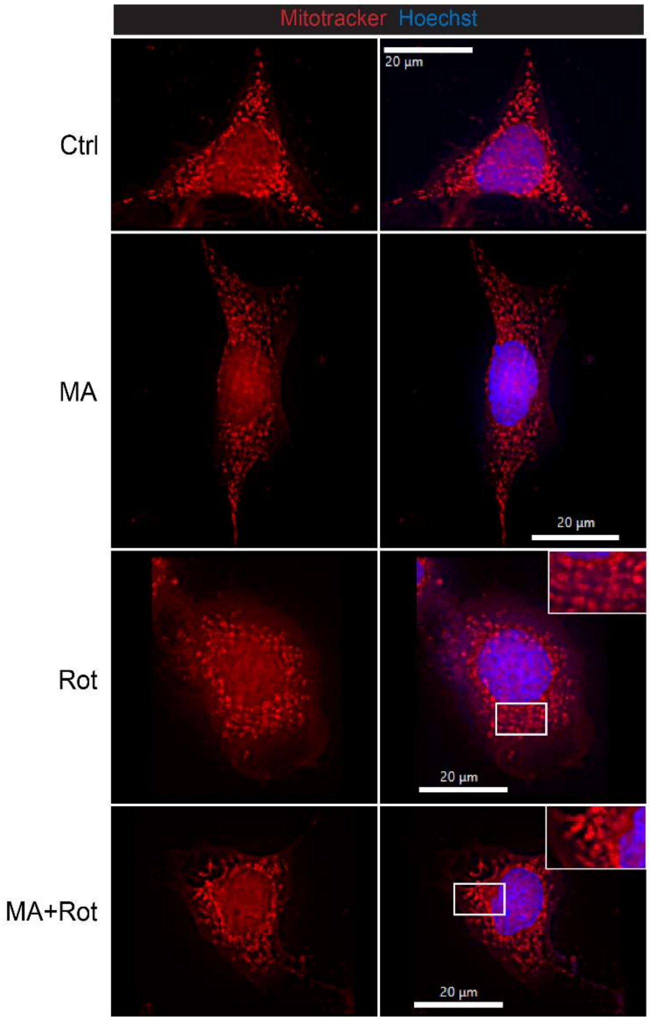


**Supp. Fig. 10 Mito-apocynin improves mitochondrial morphology in rotenone-exposed N27 DAergic neural cells**

N27 cells were incubated with 30 µM Mito-apocynin (MA), 1 µM rotenone (rot), and both Mito-apocynin and rotenone (MA+rot) for 6 h and the mitochondria were stained with Mitotracker Red dye to assess mitochondrial morphology and structural integrity. Scale bars, 20 µm.

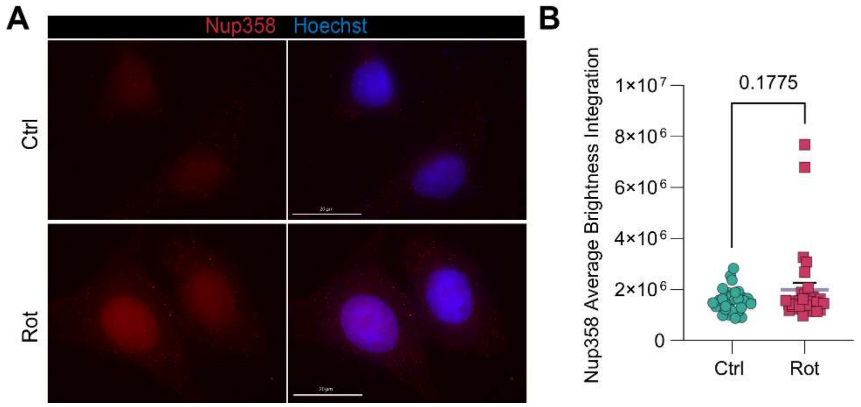


**Supp. Fig. 11 Nup358 levels in N27 DAergic neural cells.**

(A) Representative immunofluorescence images and (B) quantification of integrated fluorescence intensity of Nup358 in control and rotenone exposed N27 cells. Data from 3 independent experiments are presented as mean ± SEM and each data point represents an image. Unpaired t-tests with Welch's correction, p value shown. Scale bars, 20 μm.
